# Multiplex base editing of *BCL11A* regulatory elements to treat sickle cell disease

**DOI:** 10.1101/2024.12.13.628398

**Authors:** L. Fontana, P. Martinucci, S. Amistadi, T. Felix, M. Mombled, A. Tachtsidi, G. Corre, A. Chalumeau, G. Hardouin, J. Martin, O. Romano, M. Amendola, P. Antoniou, A Miccio

**Author notes:** equally contributed (O. Romano, M. Amendola).

## Abstract

Sickle cell disease (SCD) is a genetic anemia caused by the production of an abnormal adult hemoglobin. The clinical severity is lessened by elevated fetal hemoglobin (HbF) production in adulthood. A promising therapy is the transplantation of autologous, hematopoietic stem/progenitor cells (HSPCs) treated with CRISPR/Cas9 to downregulate the HbF repressor BCL11A via generation of double strand breaks (DSBs) in the +58-kb erythroid-specific enhancer. Here, to further enhance HbF production without increasing the mutagenic load, we targeted both +58-kb and +55-kb *BCL11A* erythroid-specific enhancers using base editors. We systematically dissected DNA motifs recognized by the key transcriptional activators within these regions and identified the critical nucleotides required for activator binding. Multiplex base editing of these residues was efficient and safe and generated no or little DSBs and genomic rearrangements. We observed substantial HbF reactivation, exceeding the levels achieved using the CRISPR/Cas9 nuclease-based strategy, thus efficiently rescuing the sickling phenotype. Multiplex base editing was efficient in long-term repopulating HSPCs and resulted in potent HbF reactivation *in vivo*. In summary, these results show that multiplex base editing of *BCL11A* erythroid-specific enhancers is a safe and potent strategy for treating sickle cell disease.

## Introduction

SCD is a highly prevalent recessive disorder caused by a single point mutation in the β-globin (*HBB*) gene, which leads to an amino acid substitution (Glu to Val) at position 6 of the β-globin chain. This sickle β-globin variant (β^s^) combines with α-chains to form sickle hemoglobin (HbS), which tends to polymerize in low oxygen conditions, causing red blood cells (RBCs) to adopt a sickle shape and lose flexibility. Clinical manifestations are due to the short lifespan of sickle RBCs (leading to anemia) and to the obstruction of small blood vessels causing multi-organ damage. Therefore, patients with SCD have a poor quality of life and reduced life expectancy ^1,2^. Beta-thalassemia is another β-hemoglobinopathy caused by mutations that reduce the production of the adult β-globin, causing severe anemia. Patients with β-hemoglobinopathies who lack a compatible donor for allogeneic HSPC transplantation can benefit from gene therapy approaches based on the transplantation of autologous, genetically modified HSPCs^3^. The severity of SCD and β-thalassemia is alleviated by the production of γ-globin chains, which compose the HbF. Gamma-globin exerts a critical anti-sickling effect and competes with β^s^-globin for incorporation into the hemoglobin tetramer, thereby reducing the formation of HbS^4^. Thus, transplantation of autologous HSPCs genetically modified to re-express HbF in their erythroid progeny is a treatment option for patients with β-hemoglobinopathies. Currently, different approaches aiming to reactivate HbF have been developed. CRISPR/Cas9 nuclease has been used to efficiently disrupt binding sites (BS) for transcriptional repressors in the γ-globin genes (*HBG1/HBG2)* promoters leading to restoration of HbF expression and rescue of the sickle phenotype^5–7^. Furthermore, CRISPR-Cas9 nuclease-mediated down-regulation of *BCL11A,* a major γ-globin repressor ^8^, was recently approved as therapy for the treatment of β-hemoglobinopathies ^9,10^. This approach targets the GATA1 activator BS within the +58-kb *BCL11A* erythroid-specific enhancer to reduce *BCL11A* expression exclusively in erythroid cells, thereby reactivating HbF production. Alternatively, targeting the ATF4 activator BS within the +55-kb *BCL11A* erythroid-specific enhancer represents another strategy to induce HbF expression in adult cells ^11^.

However, the clinical study targeting the +58-kb enhancer showed variability in the extent of HbF reactivation among individuals, relatively low levels of Hb with HbS still accounting for a large proportion of the total Hb and modest correction of ineffective erythropoiesis ^9,10^. Therefore, we hypothesize that simultaneous editing of the +58-kb and +55-kb enhancers could maximize *BCL11A* downregulation, resulting in higher and more consistent HbF levels. To avoid DSB-induced toxicity and the generation of large genomic rearrangements associated with the use of CRISPR/Cas9 nuclease ^12,13^, we used base editors (BEs) to precisely target the GATA1 and ATF4 activator BS in the +58-kb and +55-kb *BCL11A* enhancers. In particular, we exploited cytosine BE (CBEs), adenine BE (ABEs), and dual BE (DBEs) to dissect the GATA1 and ATF4 BSs in SCD HSPCs and identify the critical base conversions that induce changes in enhancers activity, *BCL11A* downregulation, and consecutively, HbF reactivation.

## Materials and Methods

### HSPC purification and culture

We obtained adult human non-mobilized CD34^+^ HSPCs from SCD patients and fetal human CD34^+^ from fetal liver of a healthy donor (HD). Adult SCD and HD fetal liver samples eligible for research purposes were obtained from the “Hôpital Necker-Enfants malades” Hospital (Paris, France). Written informed consent was obtained from all adult subjects. All experiments were performed in accordance with the Declaration of Helsinki. The study was approved by the regional investigational review board (reference: DC 2022-5364, CPP Ile-de-France II “Hôpital Necker-Enfants malades”). HSPCs were purified by immunomagnetic selection with manual magnetic cell separator (Miltenyi Biotec) after immunostaining with the CD34 MicroBead Kit (Miltenyi Biotec). Forty-eight hours before transfection, CD34^+^ cells were thawed and cultured at a concentration of 5×10^5^ cells/ml in the “HSPC medium” containing StemSpan (STEMCELL Technologies) supplemented with penicillin/streptomycin (Gibco), 250 nM StemRegenin1 (STEMCELL Technologies), and the following recombinant human cytokines (PeproTech): human stem cell factor (SCF) (300 ng/ml), FMS-like tyrosine kinase 3 ligand (Flt-3L) (300 ng/ml), thrombopoietin (TPO) (100 ng/ml), and interleukin-3 (IL-3) (60 ng/ml).

### Base editor-expressing plasmids

Plasmids used in this study include: pCMV_ABEmax_P2A_GFP (Addgene #112101), pCMV_AncBE4max_P2A_GFP (Addgene #112100), NG-ABEmax (Addgene #124163), ABE8e (Addgene #138489), SpCas9 TadDE (Addgene #193837), CBE-SpRY-OPT1 ^14^, ABE-SpRY-OPT and SpRY-ABE8e. A DNA fragment (3’UTR + poly-A) containing two copies of the 3’ untranslated region (UTR) of the *HBB* gene and a poly-A sequence of 96 adenines were purchased by Genscript. ABE-SpRY-OPT plasmid was created by inserting the 3’UTR+poly-A fragment in the pCMV-T7-SpRY-P2A-EGFP (RTW4830) (Addgene #139989) plasmid. SpRY-ABE8e plasmid was created by replacing the Cas9 sequence of the ABE8e plasmid with the SpRY-P2A-EGFP from the pCMV-T7-ABEmax(7.10)-SpRY-P2A-EGFP (RTW5025) (Addgene #140003) plasmid. Plasmids are available upon request.

### sgRNA design and production

We manually designed single guide RNAs (sgRNAs) targeting the +58-kb and +55-kb regions of *BCL11A* (**Table 1**). We used chemically modified synthetic gRNAs harboring 2′-O-methyl analogs and 3′-phosphorothioate non-hydrolyzable linkages at the first three 5′ and 3′ nucleotides (Synthego).

**Table 1.**
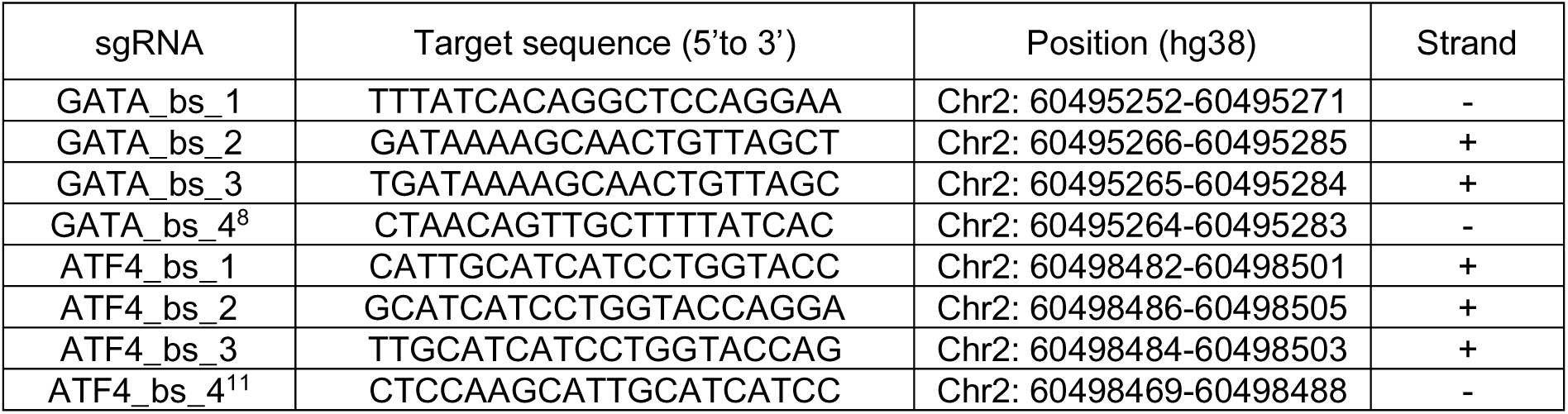
sgRNA target sequences.

### mRNA in vitro transcription

10 μg of BE-expressing plasmids were digested overnight with 20 Units of a restriction enzyme that cleaves once after the poly-A tail. The linearized plasmids were purified using a PCR purification kit (QIAGEN) and were eluted in 30 μl of DNase/RNase-free water. 1 μg of linearized plasmid was used as a template for the *in vitro* transcription (IVT) reaction (MEGAscript, Ambion). The IVT protocol was modified as follows. The GTP nucleotide solution was used at a final concentration of 3.0 mM instead of 7.5 mM and the anti-reverse cap analog N7-Methyl-3’-O-Methyl-Guanosine-5’-Triphosphate-5’-Guanosine (ARCA, Trilink) was used at a final concentration of 12.0 mM resulting in a final ratio of Cap:GTP of 4:1 that allows efficient capping of the mRNA. The incubation time for the IVT reaction was reduced to 30 minutes. mRNA was precipitated using lithium chloride and resuspended in TE buffer in a final volume that allowed to achieve a concentration of >1 μg/μl. The mRNA quality was evaluated using TapeStation 2200 (Agilent).

### RNA transfection

1×10^5^ to 2×10^5^ or 3×10^6^ CD34^+^ HSPCs per condition were transfected with 3.0 μg or 15.0 μg of the enzyme encoding mRNA, respectively, and a synthetic sgRNA at a final concentration of 2.3 μM. We used the P3 Primary Cell 4D-Nucleofector X Kit S or L (Lonza) and the CA137 program (Nucleofector 4D). Untransfected cells or cells transfected with TE buffer or with the enzyme-encoding mRNA only, or with the enzyme-encoding mRNA and a sgRNA targeting the *AAVS1* locus, served as negative controls.

### Ribonucleoprotein (RNP) transfection

RNP complexes were assembled at room temperature using a 90 μM Cas9-GFP protein and a 180 μM synthetic sgRNA (ratio Cas9:sgRNA of 1:2). CD34^+^ HSPCs (2×10^5^ cells/condition) were transfected with RNP complexes using the P3 Primary Cell 4D-Nucleofector X Kit S (Lonza) and the CA137 program (Nucleofector 4D) in the presence of a transfection enhancer (IDT). Untransfected cells or cells transfected with TE buffer served as negative controls.

### HSPC differentiation

Transfected CD34^+^ HSPCs were differentiated into mature RBCs using a three-phase erythroid differentiation protocol, as previously described ^7,15^. During the first phase (day 0 to day 6), cells were cultured in a basal erythroid medium supplemented with 100 ng/ml recombinant human SCF (PeproTech), 5 ng/ml recombinant human IL-3 (PeproTech), 3 IU/ml EPO Eprex (Janssen-Cilag) and 10^−6^ M hydrocortisone (Sigma). During the second phase (day 6 to day 9), cells were co-cultured with MS-5 stromal cells in the basal erythroid medium supplemented with 3 IU/ml EPO Eprex (Janssen-Cilag). During the third phase (day 9 to day 20), cells were co-cultured with stromal MS-5 cells in a basal erythroid medium without cytokines. Erythroid differentiation was monitored by flow cytometry analysis of CD36, CD71, CD235a, BAND3 and CD49d erythroid surface markers and of enucleated cells using the DRAQ5 double-stranded DNA dye. 7AAD was used to identify live cells.

### Colony-forming cell (CFC) assay

CD34^+^ HSPCs were plated at 1×10^3^ cells/mL concentration in a methylcellulose-based medium (Stem Cell Technologies) under conditions supporting erythroid and granulo-monocytic differentiation. BFU-E and CFU-GM colonies were counted after 14 days. Colonies were randomly picked and collected as bulk populations (containing at least 25 colonies) to evaluate base editing efficiency, globin expression by RT-qPCR and RP-HPLC, and hemoglobin expression by CE-HPLC. BFU-Es were randomly picked and collected as single colonies (around 30 colonies per sample) to evaluate base editing efficiency and globin expression by RT-qPCR.

### Genome-wide, unbiased identification of DSBs enabled by sequencing (GUIDE-seq)

Human erythroleukemia K562 cells (2.5 × 10^5^) were transfected with 500 ng of Cas9-expressing plasmid, together with 250 ng of each sgRNA–coding plasmid or an empty pUC19 vector (background control), 10 pmol of the bait double-stranded oligodeoxynucleotide (dsODN) (designed according to the original GUIDE-seq protocol^16^; and 50 ng of a pEGFP-IRES-Puro plasmid, expressing both enhanced GFP (EGFP) and the puromycin resistance genes. One day after transfection, cells were selected with puromycin (1 μg/ml) for 48 hours to enrich for transfected cells. Cells were then collected, and genomic DNA was extracted using the DNeasy Blood and Tissue Kit (Qiagen) and sheared using the Covaris S200 sonicator to an average length of 500 bp. Library preparation was performed using the original adapters and primers according to previous work ^17^.

End-repair reaction was performed using NEBNext Ultra End Repair/dA Tailing Module and adaptor ligation using NEBNext® UltraTM Ligation Module, as previously described. Amplification steps were then performed following the GUIDE-seq protocol previously described ^17^.

Libraries were sequenced with a MiSeq sequencing system (Illumina) using the Illumina MiSeq Reagent kit V2-300 cycles (paired-end sequencing; 2×150-bp). Raw sequencing data (FASTQ files) were analyzed using the GUIDE-seq computational pipeline ^16^. Identified sites were considered *bona fide* off-targets if a maximum of six mismatches against the on-target were present and if they were absent in the background control.

### Evaluation of editing efficiency

Base editing efficiency and InDel frequency were evaluated in HSPC-derived erythroid cells at the end of the first phase of differentiation and in BFU-E and CFU-GM 14 days after plating. Genomic DNA was extracted from control and edited cells using PURE LINK Genomic DNA Mini kit (LifeTechnologies), or Quick-DNA/RNA Miniprep (ZYMO Research), following manufacturers’ instructions. To evaluate base editing efficiency at sgRNA target sites, we performed PCR using primers listed in **Table 2**, followed by Sanger sequencing and EditR analysis ^18^. TIDE analysis (Tracking of InDels by Decomposition) was also performed to evaluate the percentage of InDels in edited samples ^19^.

**Table 2.**
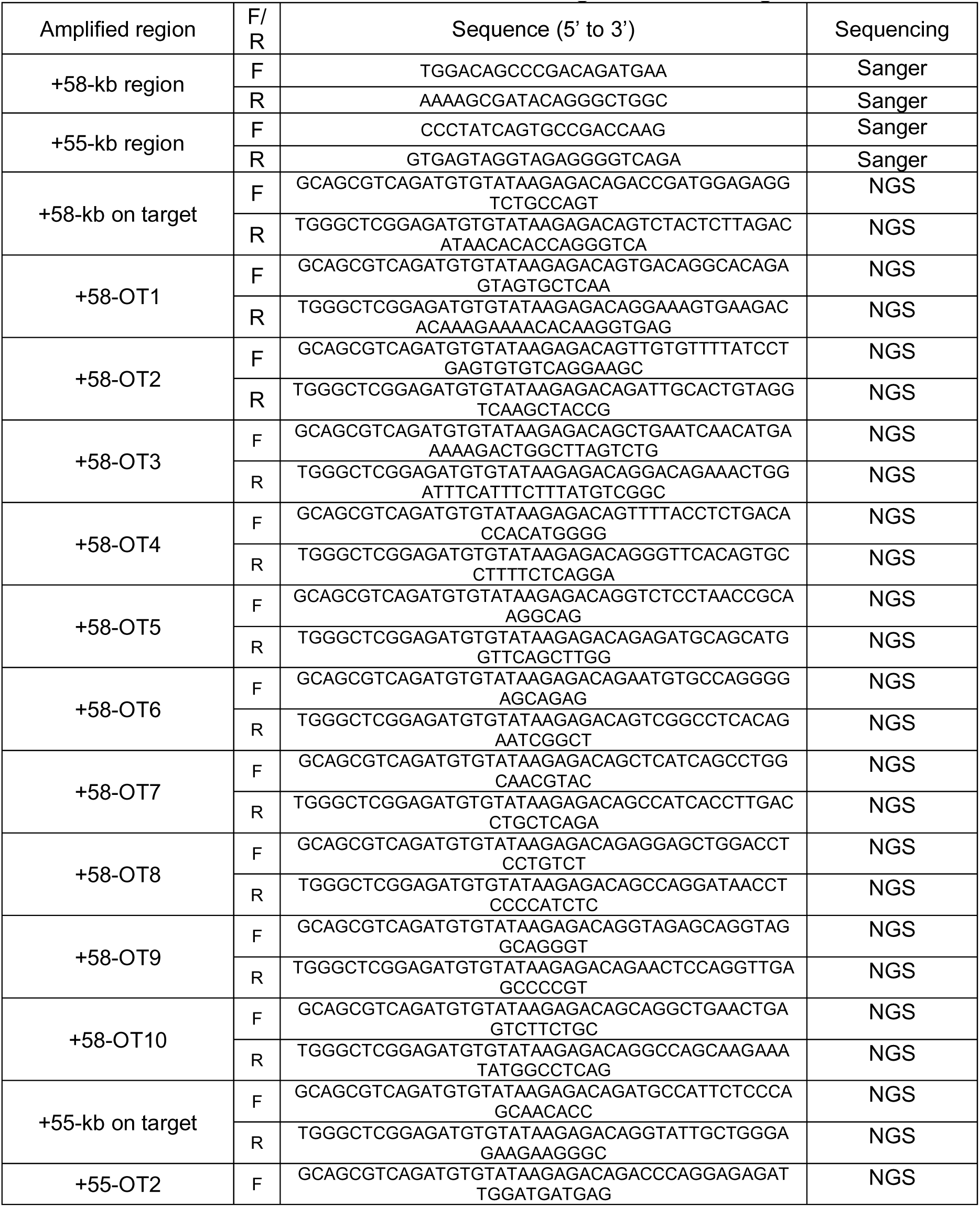
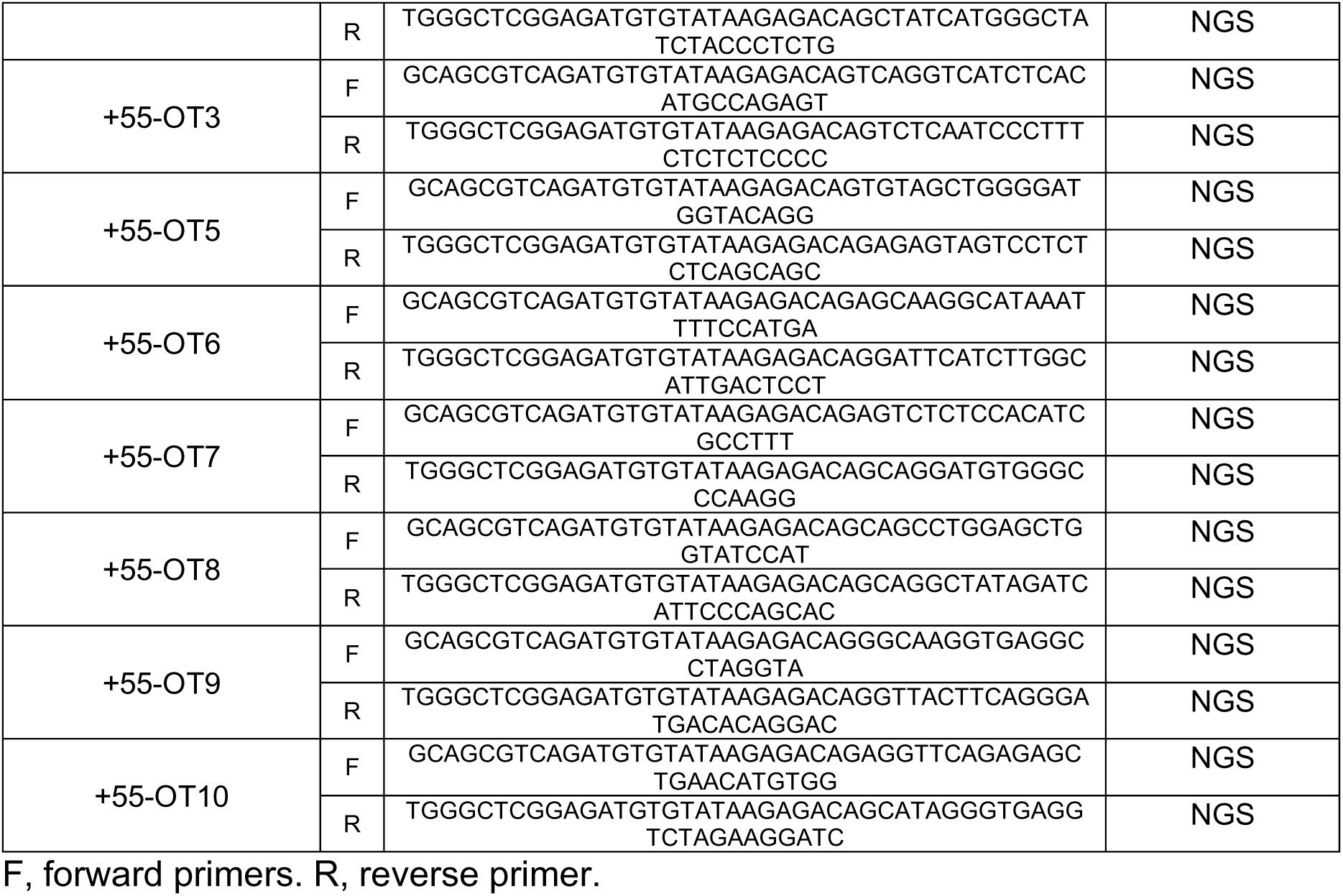
Primers used to detect on- and off-target base editing and InDel events.

On- and off-target regions were also PCR-amplified and subjected to next-generation sequencing (NGS). We selected the top 10 predicted off-targets (top 5 nominated by GUIDE-seq and top five nominated by COSMID *in silico* analysis^20^ using default parameters of mismatch tolerance of ≤3 and DNA bulge size of 1, following the COSMID algorithm criteria) and assessed editing at day 6 of culture. On-target and off-target sites were PCR-amplified using the Phusion High-Fidelity polymerase (M0530; NEB, Ipswich, MA) and primers containing specific DNA stretches (MR3 for forward primers and MR4 for reverse primers; **Table 2**) located 5’ to the sequence recognizing the off-target. Amplicons were purified using Ampure XP beads (A63881; Beckman

Coulter, Brea, CA). Illumina-compatible barcoded DNA amplicon libraries were prepared by a second PCR step using the Phusion High-Fidelity polymerase (M0530; NEB) and primers containing Unique Dual Index barcodes and annealing to MR3 and MR4 sequences. Libraries were pooled, purified using the High Pure PCR product purification kit (11732676001; Sigma-Aldrich, Saint Louis, MO), and sequenced using Illumina NovaSeq 6000 system (paired-end sequencing; 2×100-bp) to obtain a minimum of 100,000 reads per amplicon. Targeted NGS data were analyzed using CRISPResso2 ^21^.

### RT-qPCR

Total RNA was extracted from SCD HSPCs differentiated towards the erythroid lineage (day 13) using the RNeasy micro kit (QIAGEN), and from pools of BFU-E and single colonies using the Quick-DNA/RNA Miniprep kit (ZYMO Research). RNA was treated with DNase using the DNase I kit (Invitrogen), following the manufacturer’s instructions. Mature transcripts were reverse-transcribed using SuperScript First-Strand Synthesis System for RT-qPCR (Invitrogen) with oligo (dT) primers. RT-qPCR was performed using primers listed in **Table 3**, the iTaq universal SYBR Green master mix (Biorad), and the CFX384 Touch Real-Time PCR Detection System (Biorad).

**Table 3.**
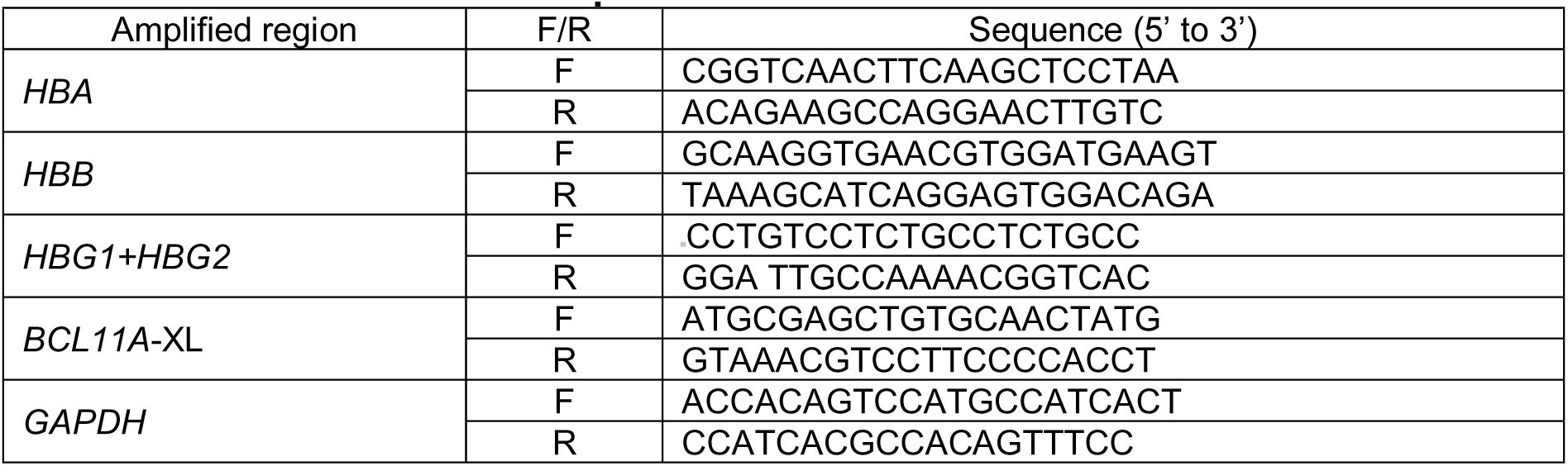
Primers used for RT-qPCR.

### Flow cytometry analysis

HSPC-derived erythroid cells were fixed with 0.05 % cold glutaraldehyde and permeabilized with 0.1 % TRITON X-100. After fixation and permeabilization, cells were stained with an antibody recognizing CD235a erythroid surface marker (PE-Cy7-conjugated anti-CD235a antibody, 563666, BD Pharmingen) and either an antibody recognizing HbF (FITC-conjugated anti-HbF antibody, clone 2D12 552829 BD), or an antibody recognizing HbS (anti-HbS antibody, H04181601, BioMedomics) followed by the staining with a secondary antibody recognizing rabbit IgG (BV421-conjugated anti-rabbit IgG, 565014, BD). Flow cytometry analysis of CD36, CD71, CD235a, BAND3 and CD49d erythroid surface markers was performed using a V450-conjugated anti-CD36 antibody (561535, BD Horizon), a FITC-conjugated anti-CD71 antibody (555536, BD Pharmingen), a PE-Cy7-conjugated anti-CD235a antibody (563666, BD Pharmingen), a PE-conjugated anti-BAND3 antibody (9439, IBGRL) and an APC-conjugated anti-CD49d antibody (559881, BD). Flow cytometry analysis of enucleated or viable cells was performed using double-stranded DNA dyes (DRAQ5, 65-0880-96, Invitrogen and 7AAD, 559925, BD, respectively). Flow cytometry analyses were performed using Gallios (Beckman coulter) flow cytometers. Data were analyzed using the FlowJo (BD Biosciences) software.

### RP-HPLC analysis of globin chains

RP-HPLC analysis was performed using a NexeraX2 SIL-30AC chromatograph and the LC Solution software (Shimadzu). A 250×4.6 mm, 3.6 μm Aeris Widepore column (Phenomenex) was used to separate globin chains by HPLC. Samples were eluted with a gradient mixture of solution A (water/acetonitrile/trifluoroacetic acid, 95:5:0.1) and solution B (water/acetonitrile/trifluoroacetic acid, 5:95:0.1). The absorbance was measured at 220 nm.

### CE-HPLC analysis of hemoglobin tetramers

Cation-exchange HPLC analysis was performed using a NexeraX2 SIL-30AC chromatograph and the LC Solution software (Shimadzu). A 2 cation-exchange column (PolyCAT A, PolyLC, Columbia, MD) was used to separate hemoglobin tetramers by HPLC. Samples were eluted with a gradient mixture of solution A (20mM bis Tris, 2mM KCN, pH=6.5) and solution B (20mM bis Tris, 2mM KCN, 250mM NaCl, pH=6.8). The absorbance was measured at 415 nm.

### Digital droplet PCR

The 3.2-kb deletion and inversion frequencies were measured by Digital Droplet PCR (ddPCR) using a primer/probe mix (Bio-rad) containing the primers and probes listed in **Table 4**. Control primers annealing to hALB (located on chr 4) were used as DNA loading control (**Table 4**). Synthetic double-stranded DNA fragments recapitulating the sequence generated after the occurrence of the 3.2-kb deletion or inversion served as positive controls.

**Table 4.**
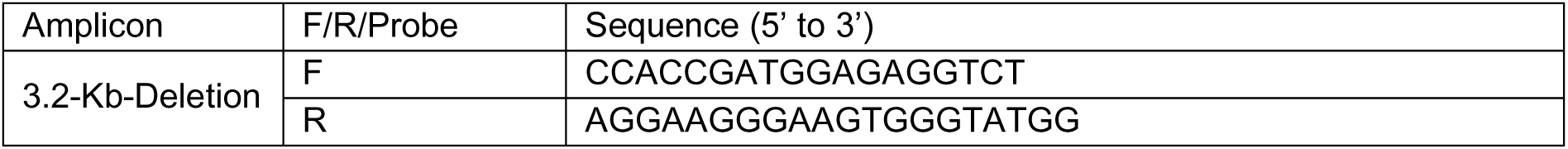
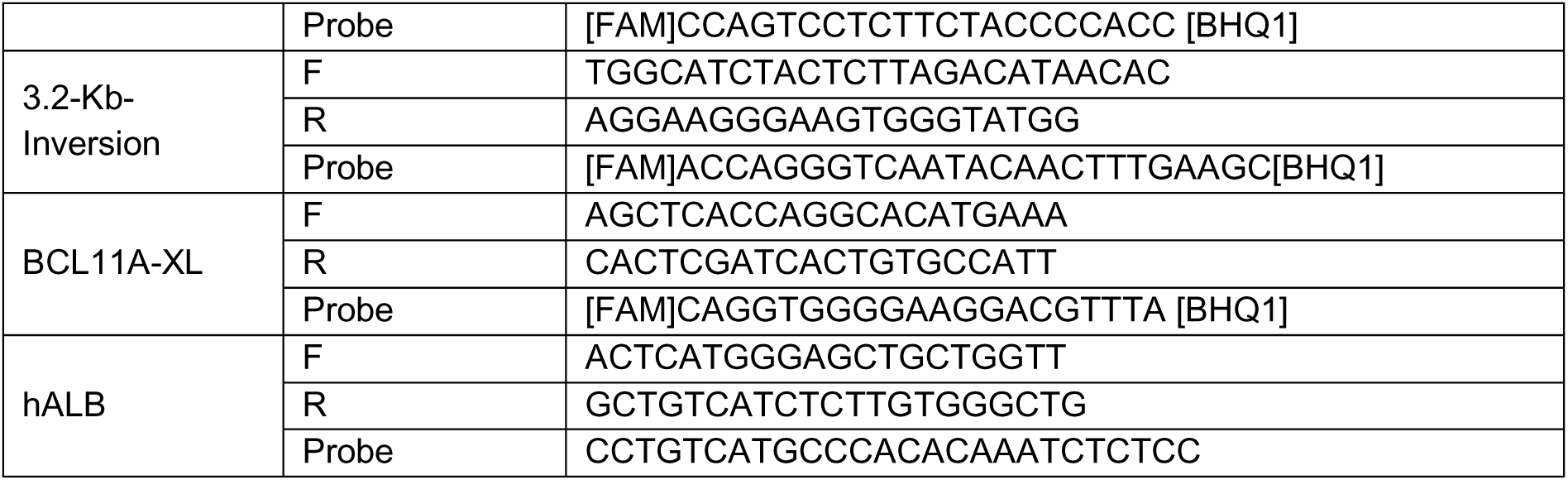
Primers used for ddPCR.

Abundance of BCL11A-XL mRNA in control and edited samples was evaluated by ddPCR. cDNA was prepared as previously described. A primer/probe mix (Bio-rad) containing the primers and probes listed in **Table 4** were used to detect BCL11A-XL. Synthetic double-stranded fragments containing BCL11A-XL sequences were used as positive controls. To detect GAPDH we used the TaqMan Gene Expression assays, VIC (Thermo).

Data were acquired through QX200 analyzer (Bio-Rad) and results were analyzed with QuantaSoftTM Analysis Pro (Bio-Rad). A positive droplet count threshold was set at 30 to allow proper calculation of copy/μL concentration through the application of the Poisson distribution.

### Sickling assay

180,000 *in vitro* differentiated RBCs (at day 19 or 20) were resuspended in CellStab (BioRad), placed in an 8-well m-Slide (Ibidi), and incubated under gradual hypoxic conditions (20% O_2_ for 20 min; 10% O_2_ for 20 min; 5% O_2_ for 20 min; 0% O_2_ for 60-180min). A time course analysis of sickling was performed in real time by video microscopy. Images were captured every 20 min using Zeiss Spinning Disk microscope and a 40x objective. Throughout the time course, images were captured and then processed with ImageJ to determine the percentage of non-sickle RBCs per field of acquisition in the total RBC population. More than 400 cells were counted per condition.

### sgRNA design for Cas9-enrichment library preparation

sgRNAs were designed to introduce cuts on complementary strands flanking the region of interest (ROI), using CHOPCHOP online design tool (https://chopchop.cbu.uib.no/) and selected for the highest predicted on-target efficiency and minimal off-target activity. The sgRNAs were assembled as a duplex from synthetic CRISPR RNAs (crRNAs) (Integrated DNA Technologies-IDT, custom designed) and trans-activating crRNAs (tracrRNAs) (IDT, catalog no. 1072532). sgRNA sequences are provided in **Table 5**. The ROI was centered on the expected cut sites of the gene editing approaches on *BCL11A* gene and has a size of 14.3 kb.

**Table 5.**
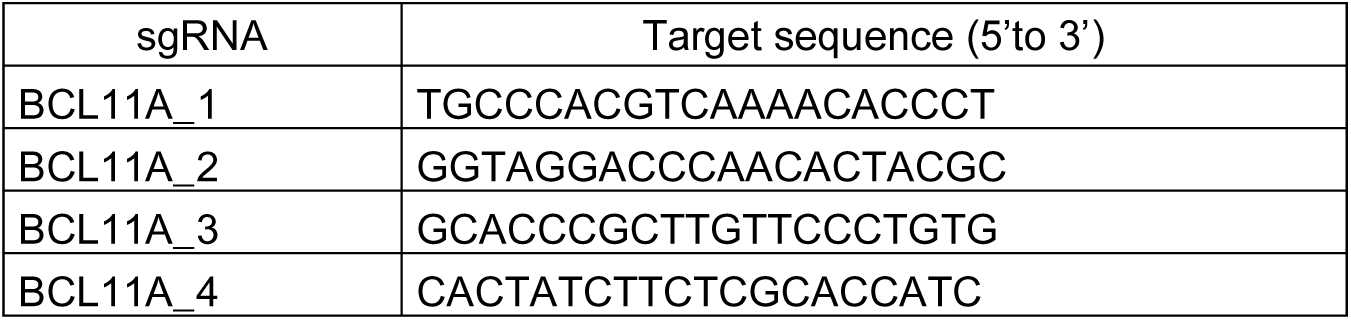
sgRNA for Cas9-enrichment library.

### Cas9-enrichment library preparation for nanopore sequencing

High molecule weight DNA was extracted from patient’s HSPC with the Nanobind CBB kit (PacBio, catalog no. 102-301-900) according to manufacturer’s instructions. DNA was size selected using Short Read Eliminator XS kit (PacBio, catalog no. 102-208-200) and quantified using the Qubit fluorometer (Thermo Fisher Scientific). 5 µg of DNA were used for the library preparation using the Cas9 Sequencing Kit (Oxford Nanopore Technologies-ONT, SQK-CS9109) and following the Cas9-mediated PCR-free enrichment protocol (version: CAS_9106_v109_revC_16Sep2020) available through ONT. Libraries were loaded onto MinION flow cells with R9.4.1 nanopores (ONT, catalog no. FLO-MIN106D). One flow cell was used per biological condition and run on GridION using MinKNOW software for 72 h.

### Bioinformatic analysis of Cas9-enrichment library from nanopore sequencing

Raw reads from Nanopore sequencing were preprocessed using cutadapt (v4.4) to remove low quality bases (q=5 threshold at 5’ and 3’ ends) and to select reads longer that 4 kb. Adaptors were then trimmed using porechop (v 0.2.4) and reads were aligned on the human reference genome (GRCh38.p13) using minimap2 (v 2.26-r1175; arguments*: -x map-ont -a -Y --secondary=no*). Reads that align at the enrichment locus (11:5220519-55121185) were re-aligned on this region using minimap2 (arguments*: -x map-ont -n 20 -I 1K -r 400,1000 -k 15 -w 15 -a -Y --secondary=no).* Finally, the depth of coverage at each position was measured using samtools (v1.17), the base composition extracted using pysamstats (v1.1.2) and the presence of INDELS detected using the variant caller sniffles (v 1.0.12; arguments*: -n -1 -r 1000 -s 1 -d 1 -l 4*). Additional processing was performed using R (v4.4.1) to prepare figures and tables. InDels were classified as small, intermediate and large according to their size ([4-50bp], [51-200bp], >200bp). The co-editing at GATA1 and ATF4 loci was assessed using mpileup output from samtools (1.21) using a 5-bp window around editing positions of the respective sgRNA (GATA_bs_3 and ATF4_bs_2).

Script & pipeline available on demand. Raw data were deposited on SRA under accession number PRJNA1192026.

### RNA-seq

Total RNA was isolated from HD HSPCs 48 h after RNA transfection using the RNeasy Kit (QIAGEN) including a DNase treatment step. Libraries were prepared using 30–50 ng of total RNA with a Watchmaker RNA kit, incorporating rRNA/Globin Polaris Depletion, following the manufacturer’s recommendations. On average 235 million paired-end reads were produced per exome library. Read quality was assessed using FastQC [version 0.11.9; https://www.bioinformatics.babraham.ac.uk/projects/fastqc/]. Adapter sequences and low-quality bases (Q < 20) were trimmed from raw reads with BBDuk [version 38.92; https://sourceforge.net/projects/bbmap/]; moreover, the first 10 nucleotides were force-trimmed for low quality. Reads shorter than 35 bp post-trimming were discarded. Trimmed reads were aligned to the human reference genome (hg38) using STAR [version 2.7.9a ^22^]. Raw gene counts were generated in R-4.1.1 using the featureCounts function of the Rsubread package [version 2.8.1 ^22,23^] and the GENCODE 44 basic gene annotation for the hg38 reference genome. Raw gene counts were normalized to counts per million mapped reads (CPM) and to fragments per kilobase of exon per million mapped reads (FPKM) using the edgeR R package [version 3.36.0 ^24^]; only genes with a CPM greater than 1 in at least 3 samples were retained for differential analysis. Differential gene expression analysis was performed using the glmQLFTest function of the edgeR R package, using the donor as a blocking variable. Genes with FDR < 0.05 and absolute log2 fold change ≥ 1 were defined as differentially expressed. Functional enrichment analysis was performed with the clusterProfiler R package [version 4.12.5 ^25^]. Fastq files and gene expression matrix from RNA-seq have been deposited in the GEO database (https://www.ncbi.nlm.nih.gov/geo/) under accession code GSE291384.

### Detection of RNA editing events by RNA-seq

RNA editing analysis was performed according to GATK Best Practices for RNA-seq variant calling (GATK v4.2.2.0). In brief, trimmed reads were two-pass aligned to the hg38 human reference genome with STAR (v. 2.7.9a)^22^ using parameters to specify the ReadGroup and output the aligned BAM file sorted by coordinate; then duplicates were marked using GATK *MarkDuplicates*. After splitting reads containing Ns in their cigar string because they span splicing sites using GATK *SplitNCigarReads*, base quality recalibration was performed using GATK *BaseRecalibrator* and *ApplyBQSR*, and the known variants collected in dbSNP155. RNA base-editing variant calling was performed using GATK *HaplotypeCaller* only on canonical (1–22, X, Y and M) chromosomes. Single-nucleotide variants (SNVs) were hard-filtered using GATK *VariantFiltration* applying suggested basic thresholds from GATK Best Practices. SNVs annotation was performed using the Variant Effect Predictor (VEP) tool from Ensembl^27^. Multiallelic variants (mainly involving repetitive sequences) were removed. Only SNVs with coverage ≥ 30 reads and genotype quality ≥ 30 were retained in each sample. To define SNVs private to treated samples (CBE, DBE and Cas9), we required a reference allele frequency ≥ 0.99 in the untreated sample (mock) at the position of the variant.

### Whole Exome Sequencing

Genomic DNA was isolated from HD HSPCs 48 h after RNA transfection using The Quik-DNA/RNA Miniprep kit (Zymo), following the manufacturer’s instructions. Exome libraries were prepared using 10 ng to 50 ng of total DNA using Twist Human Core Exome (+RefSeq) kit as recommended by the manufacturer. On average 235 million paired-end reads were produced per exome library. Read quality was evaluated using FastQC (v. 0.11.9). Adapters and low-quality tails (quality < Q20) were trimmed from raw reads with BBDuk (v. 38.92). Reads shorter than 35 bp after trimming were removed.

Variant calling was carried out according to GATK Best Practices for germline short variant discovery (GATK v4.2.2.0). In brief, FASTQ files were mapped on the hg38 human reference genome with BWA (v 0.7.17)^26^, specifying the ReadGroup. Duplicates were marked using GATK *MarkDuplicates*. Base quality recalibration was performed using GATK *BaseRecalibrator* and *ApplyBQSR*, specifying the list of target exons with a padding region of 100 bp. Variant calling was performed using GATK *HaplotypeCaller* only on canonical (1–22, X, Y and M) chromosomes. SNVs and InDels were hard-filtered using GATK *VariantFiltration* applying suggested basic thresholds from GATK Best Practices. SNVs annotation was performed using the Variant Effect Predictor (VEP) tool from Ensembl^27^. Multiallelic variants (mainly involving repetitive sequences) were removed. Only SNVs with coverage ≥ 30 reads and genotype quality ≥ 30 were retained in each sample. To define SNVs private to treated samples (CBE, DBE and Cas9), we required a reference allele frequency ≥ 0.99 in the untreated sample (mock) at the position of the variant.

Fastq files generated by WES have been deposited in the SRA database (https://www.ncbi.nlm.nih.gov/sra) under accession code PRJNA1234896.

### HSPC xenotransplantation in NBSGW mice

NOD.Cg-Kit^W-41J^Tyr^+^Prkdc^scid^Il2rg^tm1Wjl^/ThomJ (NBSGW) mice were housed in a pathogen-free facility. Control or edited mobilized CD34^+^ cells (3.5 × 10^5^ cells per mouse) were transplanted into non-irradiated NBSGW female mice of 6 to 9 weeks of age via retro-orbital sinus injection. NBSGW female mice were conditioned with busulfan (Sigma-Aldrich) injected intraperitoneally (15 mg/kg body weight) 24 hours before transplantation. Sixteen weeks after transplantation, NBSGW primary recipients were euthanized. Cells were harvested from bone marrow (BM), thymus, spleen, and blood, and stained with antibodies against the following murine and human surface markers: murine CD45 (1/50 mCD45-VioBlue; Miltenyi Biotec), human CD45 (1/50 hCD45-APCvio770; Miltenyi Biotec). BM cells were also stained with human CD3 (1/50 CD3-APC; Miltenyi Biotec), human CD14 (1/50 CD14-PECy7; BD Biosciences), human CD15 (1/50 CD15-PE; Miltenyi Biotec), human CD11b (1/100 CD11b-APC; Miltenyi Biotec), human CD19 (1/100 CD19-BV510; BD Biosciences), human CD235a (1/50 CD235a-PE; BD Biosciences), human CD71 (1/10 CD71-APC; BD Biosciences), CD36 (1/50 CD36-FITC; BD Bio-sciences), and CD34 (1/100 CD34-PE-Vio770; Miltenyi Biotec). Cells were analyzed by flow cytometry using the Novocyte analyzer (Agilent) and the FlowJo software (BD Biosciences).

Human BM CD45^+^ cells were sorted by immunomagnetic selection with CD45 MicroBeads (Miltenyi Biotec). Furthermore, BM cells were subjected to immunostaining with biotinylated antibodies that recognized the following surface markers: CD3 (dilution 1/25, clone HIT3a; BD), CD19 (dilution 1/25, clone HIB19; BD), B220 (dilution 1/50, clone RA3-6B2; BD), Ter119 (dilution 1/50, clone TER-119; BD), and mCD117 (clone 2B8; BD). BM cells were washed and incubated with 20 μL of Anti-Biotin beads (Miltenyi Biotec). After washing, the cells were magnetically purified using an LS column (Miltenyi Biotec) according to the manufacturer’s instructions. Cells from the positive fraction were immuno-stained with the following antibodies: CD19-PE (dilution 1/20, BD), streptavidin (SA)-APC (dilution 1/20 BD) and hCD45-BV510 (dilution 1/100, BD). The hCD45^high^/CD19^high^ were sorted using MA900 cell sorter and subjected to RT-qPCR analysis. Cells from the negative fraction were immuno-stained with the following antibodies: CD235a-PE (dilution 1/5000, BD) and hCD45-BV510 (dilution 1/100, BD). The hCD45^low/−^/CD235a^high^ cells were sorted using the MA900 cell sorter (Sony Biotechnology, San Jose, CA) and subjected to flow cytometry, reverse phase high-performance liquid chromatography (RP-HPLC) and RT-qPCR analysis.

All experiments and procedures were performed in compliance with the French Ministry of Agriculture’s regulations on animal experiments and were approved by the regional Animal Care and Use Committee (APAFIS#2019061312202425_v4). Mice were housed in a temperature-(20◦C-22◦C) and humidity (40%-50%)-controlled environment with 12:12 hour light-dark cycle and fed *ad libitum* a standard diet.

## Results

### Base editing disrupts the +58-kb and +55-kb BCL11A enhancers without affecting SCD HSPC viability and differentiation

To disrupt the *BCL11A* erythroid enhancers, we targeted the +58-kb and +55-kb regions in human SCD HSPCs by inserting point mutations at the GATA1 and ATF4 BS, respectively (**Figure 1A**). SCD HSPCs were transfected with various combinations of *in vitro* transcribed BE mRNAs and chemically modified sgRNAs, and afterwards subjected to erythroid differentiation (**Figure 1B**). We evaluated the base editing efficiency in erythroblasts derived from transfected HSPCs. At both the +58-kb and +55-kb regions, different combinations of CBEs/ABEs and sgRNAs led to the generation of several editing profiles. In the +58-kb region, CBEs created the +58 CBEI profile (<u>G</u>T<u>G</u>ATAAA; targeted nucleotides are underlined; 55.6% ± 4.7) and ABEs created the +58 ABEI (G<u>T</u>GA<u>T</u>AAA; 28.4% ± 6.1), +58 ABEII (GTGAT<u>AAA</u>; 19.0% ± 3.3) and +58 ABEIII profiles (GTGAT<u>A</u>AA; 14% ± 2.0; **Figure 1C**). In the +55-kb region, CBEs led to the +55 CBEI (TTG<u>C</u>AT<u>C</u>AT<u>CC</u>; 47.0% ± 7.0) and +55 CBEII profiles (TTGCAT<u>C</u>AT<u>CC</u>; 46.7% ± 3.7), while ABEs generated the +55 ABEI (TTGC<u>A</u>TCATCC; 71% ± 3.5) and the +55 ABEII profiles (TTGC<u>A</u>TC<u>A</u>TCC; 87.0% ± 5.1; **Figure 1D**). To compare base editing and Cas9 nuclease strategies, we transfected Cas9 nuclease RNP complexes targeting either the +58-kb^8^ or the +55-kb^11^ region and disrupted the GATA1 and ATF4 BS, through InDel generation (75.8% ± 3.0 and 82.3% ± 4.1, respectively; **Figure 1E-F**). Low or no InDels were detected in base edited samples, confirming the DSB-low nature of BEs (**Figure 1E-F**). Transfected SCD HSPCs were plated on a semi-solid medium that allows erythroid and granulocyte/monocyte differentiation at clonal level (CFC assay). No differences in the frequency of erythroid (BFU-E) and granulocyte/monocyte (CFU-GM) colonies were detected between control and edited samples, demonstrating that our therapeutic strategy does not affect progenitors’ viability and differentiation towards the erythroid and granulomonocytic lineages (**Figure 1G**). The base editing efficiency and InDel profiles in pools of BFU-E and CFU-GM were similar to those measured in liquid erythroid cultures (**Supplementary Figure 1A-H)**. In conclusion, we efficiently targeted the GATA1 and ATF4 BSs at the +58-kb and +55-kb *BCL11A* enhancer regions with base editing strategies without affecting HSPCs viability and differentiation potential. This approach allowed for precise modifications of alternative bases, resulting in significantly fewer DSBs compared to Cas9 nuclease.

**Figure 1.**
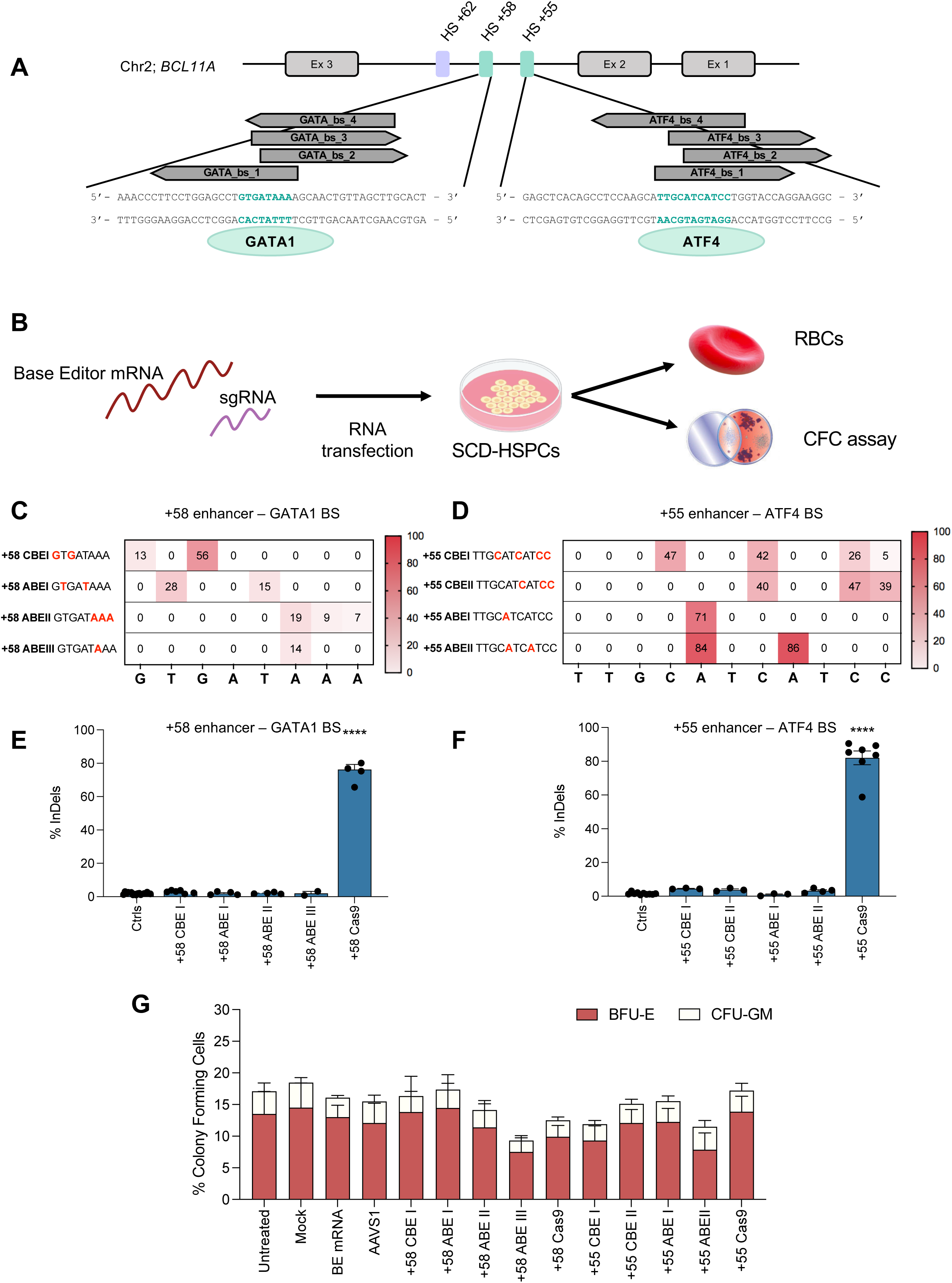
Base editing of the erythroid-specific *BCL11A* enhancers in SCD HPSC-derived erythroblasts. **A.** Schematic representation of part of the *BCL11A* gene on chromosome 2, depicting exons 2, 3 and 4 (Ex 2, 3 and 4) and the DNaseI hypersensitive sites (HS) +62-kb, +58-kb and +55-kb. The sequences of the +58-kb and +55-kb regions of the erythroid-specific *BCL11A* enhancers are depicted. GATA1 and ATF4 transcriptional activator binding sites (BSs) are highlighted in green and bold. Target sgRNAs used in combination with base editing enzymes are indicated with arrows and aligned with the DNA sequence that they bind to in a stranded-oriented way. Green ovals indicate the transcriptional activators GATA1 and ATF4. **B.** Experimental protocol used for base editing experiments in SCD HSPCs. A BE mRNA and a sgRNA were co-transfected in SCD HSPCs. Cells were differentiated into mature RBCs using a three-phase erythroid differentiation protocol or plated in methylcellulose-containing medium under conditions supporting erythroid (BFU-E) and granulo-monocytic (CFU-GM) differentiation. **C-D**. C-G to T-A or A-T to G-C base-editing efficiency calculated by the EditR software in samples subjected to Sanger sequencing in erythroblasts derived from SCD HSPCs edited in the +58-kb (**C**) or +55-kb (**D**) regions. Different editing profile were generated through the combination of different BEs and sgRNA. GATA_bs_1 sgRNA was used in combination with CBE-SpRY or AncBE4max and ABEmax-SpRY to generate the +58 CBEI and +58 ABEI profile, respectively. GATA_bs_2 sgRNA was coupled with NG-ABEmax and ABEmax-SpRY to generate +58ABEII profile while GATA_bs_3 sgRNA was combined with ABEmax-SpRY generating the +58 ABEIII profile. CBE-SpRY and AncBE4max coupled with ATF4_bs_1 or ATF4_bs_2 sgRNAs led to +55 CBEI and +55 CBEII profiles, respectively. ATF4_bs_1 sgRNA or ATF4_bs_3 combined with ABEmax-SpRY led to +55 ABEI profile and ATF4_bs_2 sgRNA combined with ABE8e-SpRY led to +55 ABEII profile. In the results, we reported the highest editing efficiency achieved in the editing window. Data are expressed as mean (n = 2 to 6 biologically independent experiments, 2 to 5 donors). **E-F**. Frequency of InDels, measured by TIDE analysis, in samples subjected to Sanger sequencing in erythroblasts derived from SCD HSPCs edited in the +58-kb (**E**) or +55-kb (**F**) regions. Data are expressed as mean ± SEM (n = 2 to 6 biologically independent experiments, 2 to 5 donors). ****P≤ 0.0001 (One-way ANOVA. Comparison of controls vs edited samples. No asterisk (*) = not significant. **G**. CFC frequency for controls (untreated, or transfected with TE buffer, or transfected with a BE mRNA only, or transfected with a BE mRNA and a sgRNA targeting the unrelated *AAVS1* locus) and edited samples. Data are expressed as mean ± SEM (n = 7 biologically independent experiments, 7 donors) (Two-way ANOVA with Dunnet correction for multiple comparisons; not significant).

### HbF reactivation after base editing of BCL11A enhancers in SCD HSPCs

We differentiated transfected SCD HSPCs in liquid culture towards the erythroid lineage to produce mature RBCs. Flow cytometry analysis of enucleation and early and late erythroid markers showed no difference between control and edited groups, indicating no impairment of SCD HSPC erythroid differentiation upon targeting of the GATA1 and the ATF4 BS in the *BCL11A* erythroid enhancers (**Supplementary Figure 2A-E**). We assessed *HBG* reactivation at RNA and protein levels by RT-qPCR, RP-HPLC and CE-HPLC (**Figure 2A-C**). Regarding the +58-kb region, CBE-treated cells bearing the +58 CBEI profile, expressed the highest HbF levels among base edited groups and compared to the control groups. Notably, the levels of HbF obtained in cells carrying the +58 CBEI profile were only modestly lower compared to the +58 Cas9 profile, despite the significantly lower editing efficiency (56.0% ± 4.7 and 76% ± 3.0 in +58 CBEI and +58 Cas9, respectively). ABE-treated samples exhibited low HbF levels that were consistent with limited editing efficiency (**Figure 2A-C**). Interestingly, cells carrying the +58 ABEIII profile showed HbF levels similar to those achieved with ABEI and ABEII, despite having the lowest editing, suggesting that this base conversion is potentially highly productive in terms of HbF reactivation (**Figure 2A-C**).

**Figure 2.**
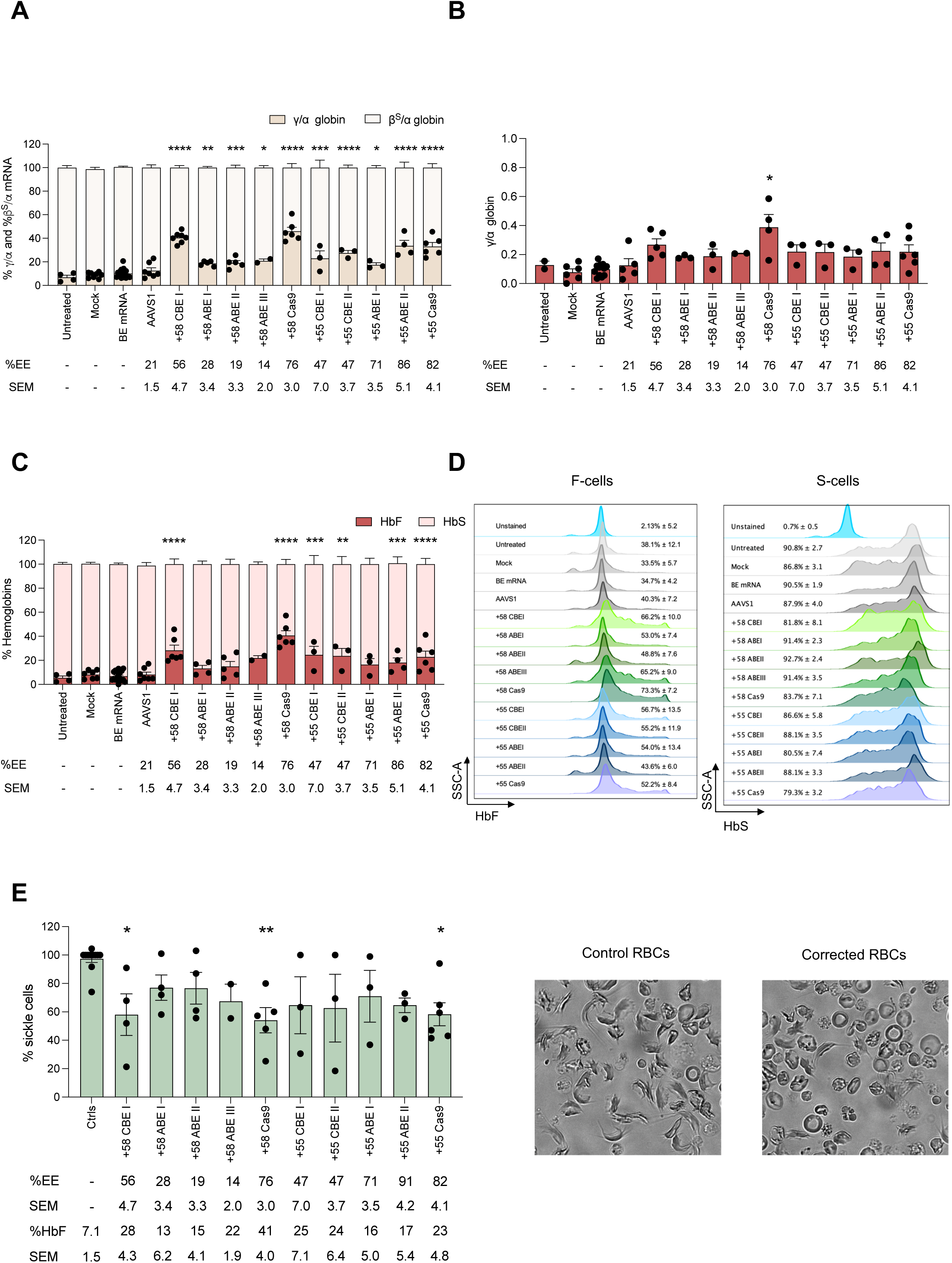
HbF reactivation and correction of the sickle phenotype in edited SCD HPSC-derived erythroblasts. **A.** RT-qPCR analysis of γ (^G^γ + ^A^γ)- and β^S^-globin mRNA in control and edited SCD patient erythroblasts at day 13 of erythroid differentiation. γ- and β^S^-globin mRNA expression was normalized to α-globin mRNA and expressed as a percentage of the γ- + β^S^-globin mRNA. The editing efficiency (EE) ± SEM is indicated for each sample in the lower part of the panel. Data are expressed as mean ± SEM (n = 2 to 6 biologically independent experiments, 2 to 5 donors). *P ≤ 0.05; **P ≤ 0.01; ***P ≤ 0.001; ****P ≤ 0.0001 (two-way ANOVA with Dunnett correction for multiple comparisons. Statistical significance between mock and edited samples is depicted in the graph). **B.** Expression of γ (^G^γ + ^A^γ)-globin chains measured by RP-HPLC in SCD RBCs derived from control and edited SCD HSPCs. γ-globin expression was normalized to α-globin. The EE ± SEM is indicated for each sample in the lower part of the panel. Data are expressed as mean ± SEM (n = 2 to 6 biologically independent experiments, 2 to 5 donors). *P ≤ 0.01 (one-way ANOVA with Dunnett correction for multiple comparisons. Statistical significance between mock and edited samples is depicted in the graph). **C.** Analysis of HbF and HbS expression measured by cation-exchange HPLC in RBCs derived from control and edited SCD HSPCs. We calculated the percentage of each Hb type over the total Hb tetramer. The EE ± SEM is indicated for each sample in the lower part of the panel. Data are expressed as mean ± SEM (n = 2 to 6 biologically independent experiments, 2 to 5 donors). *P ≤ 0.05; **P ≤ 0.01; ***P ≤ 0.001; ****P ≤ 0.0001 (two-way ANOVA with Dunnett correction for multiple comparisons. Statistical significance between mock and edited samples is depicted in the graph). **D.** Representative flow cytometry histograms showing the percentage of HbF-expressing cells (F-cells) and HbS-expressing cells (S-cells) in CD235a^+^ population for unstained (CD235a stained only), control and edited samples. Data expressed as mean ± SEM of F-cells and S-cells (n = 2 to 4 biologically independent experiments, 2 to 5 donors). **E.** Frequency of sickling cells upon O_2_ deprivation in control and edited samples after normalization to the mock. The EE ± SEM is indicated for each sample in the lower part of the panel. Representative photomicrographs of SCD patient RBCs under hypoxia conditions are shown. Data are expressed mean ± SEM (n = 2 to 6 biologically independent experiments, 2 to 5 donors). *P ≤ 0.05; **P ≤ 0.01 (one-way ANOVA with Dunnett correction for multiple comparisons. Statistical significance between mock and edited samples is depicted in the graph).

Concerning the +55-kb region, CBE- and ABE-treated samples showed HbF reactivation, with an extent similar to Cas9-treated samples even though the CBE editing frequency was around two-fold lower compared to ABE and Cas9 (**Figure 2A-C**).

Similar results in term of HbF reactivation were observed also in pools of BFU-E (**Supplementary Figure 1I**). Flow cytometry analysis of F- and S-cells evidenced similar results in terms of HbF-expressing cells with a concomitant slight reduction of HbS-expressing cells (**Figure 2D**). Finally, a sickling assay was performed in control and edited samples. Edited samples showed decreased frequencies of sickle cells, that were in accordance with the levels of HbF (**Figure 2E**), with +58 CBEI, +58 Cas9, + and +55 Cas9 profiles causing the highest reduction in the frequency of sickle cells (**Figure 2E**). Overall, these data showed that base editing-mediated disruption of the GATA1 or ATF4 activator BS in the +58-kb or +55-kb regions, respectively, led to HbF reactivation, which, however, was at best only approaching that achieved with the approved strategy using Cas9 to target the +58-kb region. Moreover, phenotypic rescue (evaluated by performing a sickling assay) was incomplete and inconsistent across different donors for all editing profiles, including +58 Cas9, indicating the need for further optimization of the approach. We hypothesize that this can be accomplished by further increasing HbF expression.

### Dissecting GATA1 and ATF4 binding motifs to achieve robust HbF reactivation

To precisely evaluate the potency of the different editing profiles, we correlated γ-globin expression and editing efficiency at the clonal level in BFU-Es derived from edited SCD HSPCs. Single colonies were analyzed for *HBG1/2* mRNA expression, base editing efficiency or InDel frequency (**Figure 3A and B**). We observed a positive correlation between editing efficiency and γ-globin reactivation in all the groups. For the +58-kb enhancer, the generation of the +58 ABEIII profile was significantly the most potent profile in terms of γ-globin reactivation, outperforming even the +58 Cas9 profile (**Figure 3A**). To increase the editing efficiency in the +58 ABEIII profile, we used a more efficient enzyme, ABE8e^28^ instead of the first generation ABEmax. However, due to a wider editing window, we were not able to generate the +58 ABEIII profile, but instead, we recreated the +58 ABEII profile with an editing efficiency of 80.5% ± 0.7 that was associated with a higher Indel frequency of 4.4% ± 2.8 (**Figure 3 C and D**). Conversely, higher editing efficiency obtained for the +58 ABEI and +58 ABEII profiles led to the generation of bi-allelic colonies, making these profiles more suitable for future experiments (**Figure 3A**). Given the higher number of bi-allelic colonies and the better correlation between editing frequency and *HBG1/2* levels, the +58 ABEII profile was selected for further testing. The results emphasize the importance of specific bases in the GATA1 BS motif for GATA1 binding and γ-globin regulation. Targeting the A6 position (by generating the +58 ABEIII profile: GTGAT<u>A</u>AA) significantly increased γ-globin expression, indicating its pivotal role. However, simultaneous targeting of A6 to A8 (+58 ABEII) reduced γ-globin expression. Additionally, T>C conversions at T2 and T5 (+58 ABEI) were less effective in reactivating HbF than the generation of the ABEIII profile, indicating a lower impact on GATA1 binding capacity (**Figure 3A**). Finally, the activity of the +58 CBEI profile mirrored that of the +58 Cas9 profile, leading to a high γ-globin reactivation. This confirmed previously reported observations of the importance of G in position 3 for GATA1 binding^29^; thus, the +58 CBEI profile was also chosen for further testing (**Figure 3A**).

**Figure 3.**
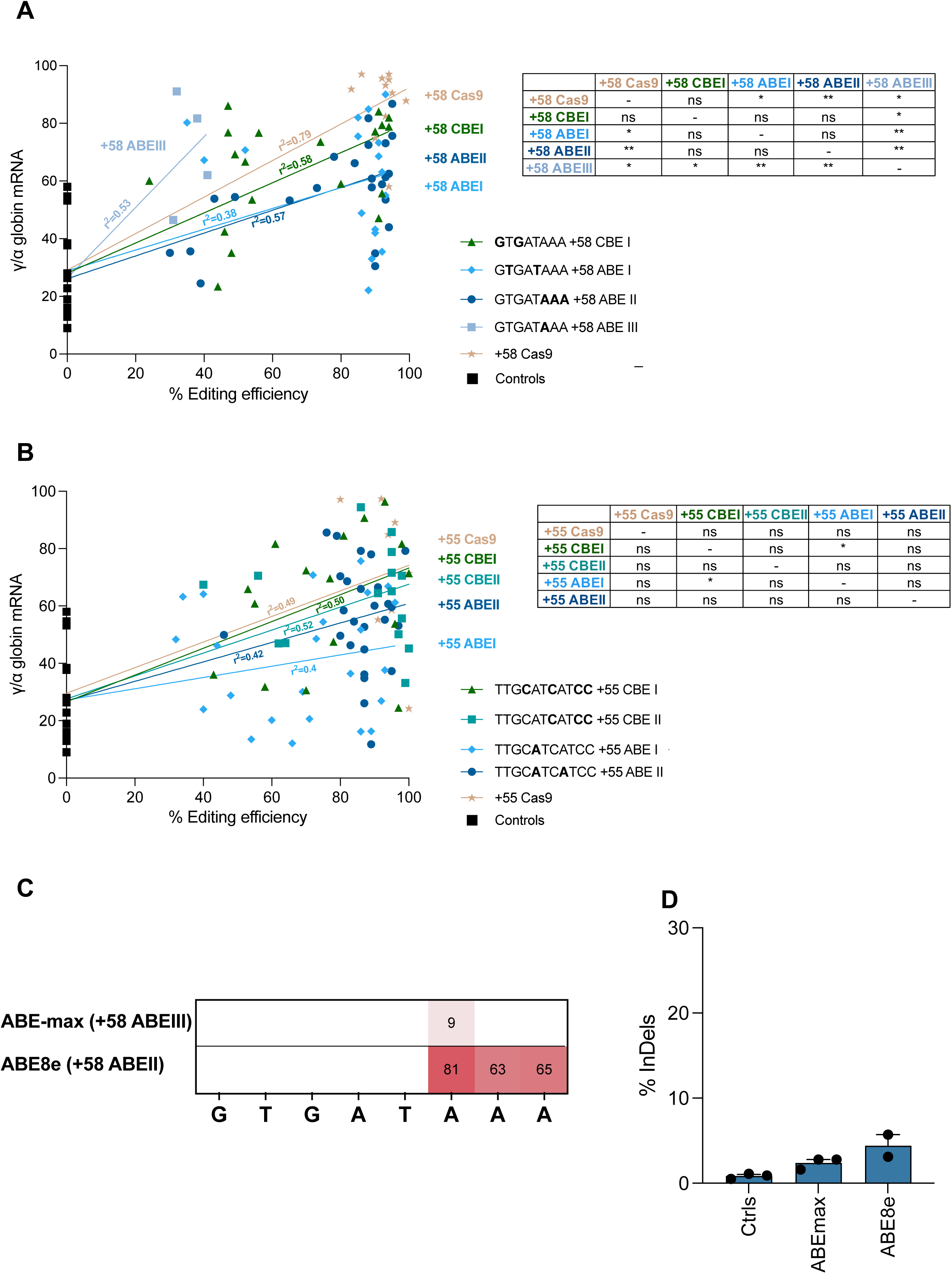
Gamma-globin reactivation in single BFU-E colonies derived from edited SCD HSPCs. **A-B** Correlation between *HBG* mRNA relative expression and editing efficiency in single BFU-E colonies differentiated from SCD HSPCs edited in the +58-kb (**A**) or +55-kb (**B**) regions (1 donor). Different editing profiles were generated through the combination of different BE and sgRNA. Different editing profiles were generated through the combination of different BE and sgRNA similar to Figure 1C-D. mRNA expression was measured by RT-qPCR analysis of γ (^G^γ + ^A^γ)- and β^S^-globin mRNA in control and edited BFU-E. γ- and β^S^-globin mRNA expression was normalized to α-globin mRNA and expressed as a percentage of the γ-+ β^S^-globin mRNA. Base-editing efficiency was calculated using EditR and Cas9-editing efficiency (InDels) was calculated using TIDE in samples subjected to Sanger sequencing. BFU-Es transfected with TE buffer, or transfected with a BE mRNA only, or transfected with a BE mRNA and a sgRNA targeting the unrelated AAVS1 locus were used as negative controls. Statistical significances are reported in the tables *P ≤ 0.05; **P ≤ 0.01 (Multiple t test). **C.** A-T to G-C base-editing efficiency calculated by the EditR software in samples subjected to Sanger sequencing in pools of BFU-E derived from SCD HSPCs edited in the +58-kb region. Data are expressed as mean ± SEM (n = 2 to 3 biologically independent experiments, 2 to 3 donors). **D.** Frequency of InDels, measured by TIDE analysis, in samples subjected to Sanger sequencing in pools of BFU-E colonies derived from SCD HSPCs edited in the +58-kb. The GATA_bs_2 sgRNA was combined with NG-ABE-max and NG-ABE8e. Data are expressed as mean ± SEM (n = 2 to 3 biologically independent experiments, 2 to 3 donors). **P ≤ 0.01 (One-way ANOVA comparison. Statistical significance between mock and edited samples is depicted in the graph).

With regards to the +55-kb enhancer, all the profiles exhibited comparable γ-globin reactivation (with the exception of the +55 ABEI showing the lowest *HBG1/2* levels) but we selected +55 CBEII and +55 ABEII profiles that showed a better correlation between editing frequency and *HBG1/2* levels **(Figure 3B**). These data suggest that insertion of C>T conversion at C4 of the binding motif to generate +55 CBEI profile (TTG<u>C</u>AT<u>C</u>AT<u>CC</u>) does not further impact the binding capacity of ATF4 compared to targeting only the last 3 Cs in the +55 CBEI profile (TTGCAT<u>C</u>AT<u>CC</u>). On the other hand, reduced γ-globin levels were associated with the +55 ABEI and +55 ABEII profiles, suggesting that A>G mutations (and in particular targeting of only the A5 of the ATF4 motif in the +55 ABEI profile) does not strongly impact ATF4 binding.

### Enhanced γ-globin reactivation through multiplex editing of +58 and +55 enhancers

We hypothesized that simultaneous editing of the +58-kb and +55-kb enhancers of SCD HSPCs could lead to a stronger reduction of *BCL11A* expression, and consequently greater HbF induction compared to the editing of a single enhancer alone. We transfected SCD HSPCs with both base editors and sgRNAs, aiming at creating the most potent editing profiles simultaneously (**Figure 3A and B**). In erythroid liquid cultures, modestly reduced editing efficiencies were observed when inserting the combination of +58 ABEII and +55 ABEII profiles compared to samples transfected with the individual ABE/sgRNA combinations (70% ± 18.1 and 52.0% ± 11.5 achieved for +58 ABEII/+55 ABEII profiles at GATA1 and ATF4 BS respectively, compared to 77.7 % ± 7.6 and 60.0% ± 5.7 obtained for +58 ABEII and +55 ABEII single editing profiles, respectively; **Figure 4A and B**). We then used CBEs to insert the most potent CBE profiles, +58 CBEI and +55 CBEII, reaching editing efficiencies similar to those achieved by single editing (61.3% ± 15.5 and 54.3% ± 17.0, respectively, for the combined +58 CBEI/+55 CBEII profile compared to 60.7% ± 14.5 and 45.3% ± 22.8 for +58 CBEI and +55 CBEII individual profiles, respectively; **Figure 4A and B**). We also used a recently developed, dual function, cytosine and adenine BE (TadDE; Neugebauer et al., 2022), to simultaneously introduce C>T and A>G point mutations at the GATA1 and ATF4 BSs, generating two additional profiles (+58 DBEI: G<u>TG</u>ATAAA; +55 DBEI: TTGCAT<u>CA</u>T<u>CC</u>), reaching high editing efficiencies upon single or multiplex editing (65.3% ± 22.1 and 57.7% ± 5.0 for +58 DBEI and +55 DBEI individual profiles, and 67.0% ± 13.0 and 49% ± 7.2 for the combined +58 DBEI/+55 DBEI profile; **Figure 4A and B**). Importantly, base-edited samples showed no or few InDels (**Figure 4C and D**). In parallel, we used Cas9 nuclease to simultaneously target both enhancers. The efficiency of concomitant generation of the +58 and +55 Cas9 profiles was similar to that obtained upon single Cas9 editing (41.9% ± 9.6 and 40.6% ± 15.4, when targeting the GATA1 and ATF4 BS simultaneously and 49.6% ± 9.6 and 44.7% ± 21.1 for individual +58 Cas9 and +55 Cas9 profiles, respectively; **Figure 4C and D**). This approach potentially generates a 3.2-kb deletion or inversion that disrupts the targeted GATA1 and ATF4 BSs and removes additional BSs for transcriptional activators (**Figure 4E**). ddPCR showed 33.1% ± 8.1 of deletion and 10.4% ± 2.3 of inversion for the +58/+55 Cas9 profile (**Figure 4F**). On the other hand, no deletion or inversion was detected with the +58 ABEII/+55 ABEII profile, and only 0.5% ± 0.4 and 1.1% ± 0.9 of deletion were detected for +58 CBEI/+55 CBEII and +58 DBEI/+55 DBEI profiles, respectively (**Figure 4F**). A CFC assay demonstrated no difference in terms of viability and clonogenic potential between control, single- and multiplex-edited SCD HSPCs (**Figure 4G**). The frequency of base editing, InDel and 3.2-kb deletion/inversion in the pools of BFU-E and CFU-GM populations were similar to those measured in the liquid erythroid cultures (**Supplementary Figure 3C-L**).

**Figure 4.**
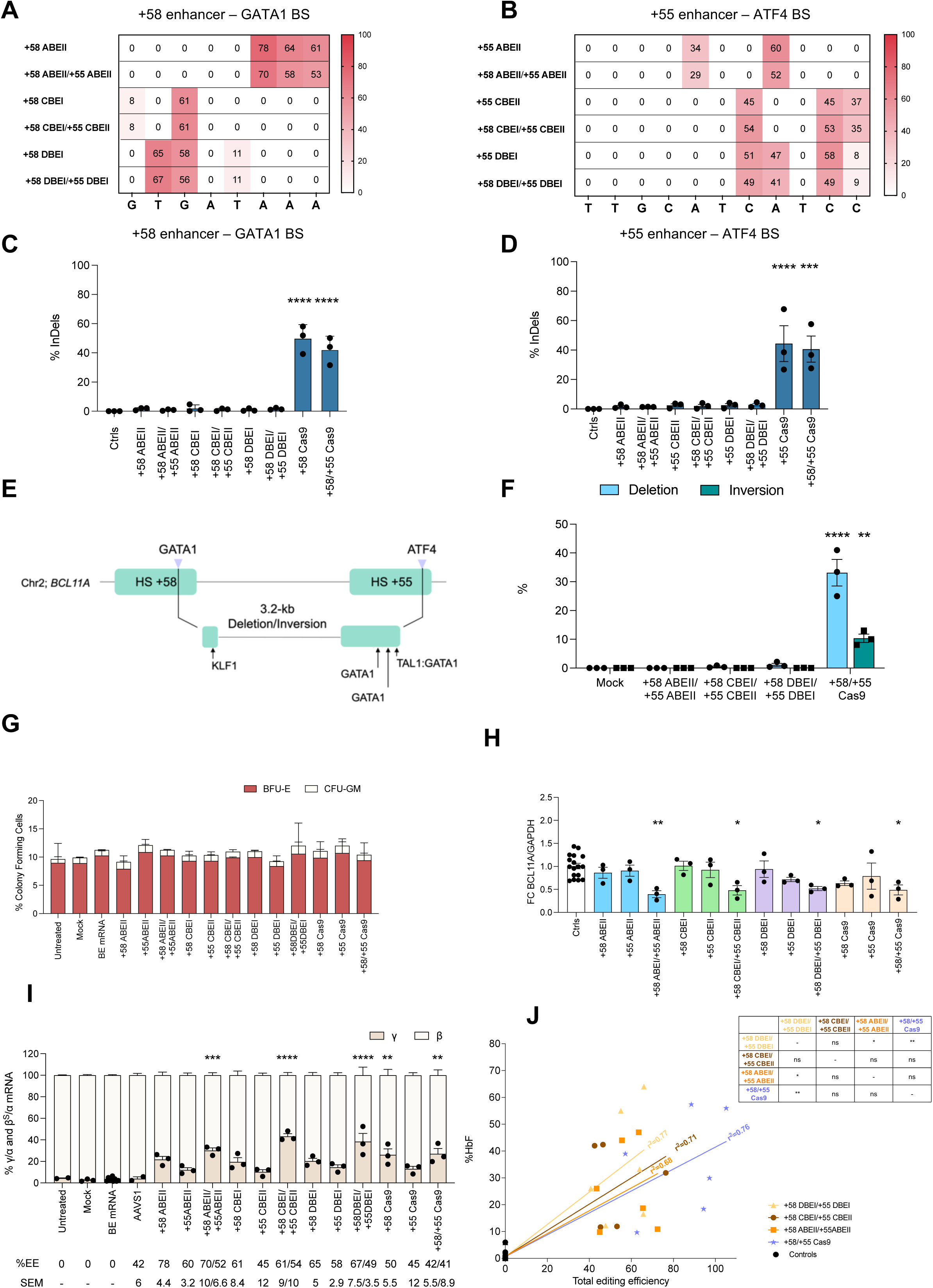
Multiplex base editing of the erythroid-specific *BCL11A* enhancers in SCD HPSC-derived erythroblasts. **A-B**. C-G to T-A or/and A-T to G-C base-editing efficiency in the +58-kb (**A**) or +55-kb **(B**) regions calculated by the EditR software in sample subjected to Sanger sequencing in erythroblasts derived from SCD HSPCs edited at either the +58-kb or the +55-kb region, or simultaneously edited at both the +58-kb and +55-kb regions. GATA_bs_2 or/and ATF4_bs_2 sgRNAs were combined with NG-ABE8e to generate +58ABEII, +55ABEII, and +58ABEII/+55ABEII profiles. GATA_bs_1 or/and ATF4_bs_2 sgRNAs combined with AncBE4max, generated the +58CBEI, +55CBEII, and +58CBEI/+55 CBEII profiles. Finally, GATA_bs_1 or/and ATF4_bs_2 sgRNAs combined with TadDE, generated the +58DBEI, +55DBEI, and +58DBEI/+55 DBEI profiles. Data are expressed as mean (n = 3 biologically independent experiments, 3 donors). **C-D**. Frequency of InDels in the +58-kb (**C**) or +55-kb (**D**) regions measured by TIDE analysis, in samples subjected to Sanger sequencing in erythroblasts derived from SCD HSPCs edited at either the +58-kb or the +55-kb regions or simultaneously edited at both the +58-kb and the +55-kb regions. Data are expressed as mean ± SEM (n = 3 biologically independent experiments, 3 donors). ****P ≤ 0.0001; *** ≤ 0.001 (One-way ANOVA. Comparison of controls vs edited samples) **E.** Schematic representation of the generation of a 3.2-kb deletion/inversion following the simultaneous targeting of GATA1 BS in the +58-kb region and ATF4 BS in the +55-kb region (highlighted with violet arrows). Black arrows indicate the location of BSs for additional transcriptional activators. **F.** Frequency of the 3.2-kb deletion/inversion, measured by ddPCR, for samples simultaneously edited at the +58-kb and +55-kb regions. Data are expressed as mean ± SEM (n = 3 biologically independent experiments, 3 donors). **P ≤ 0.01; ****P ≤ 0.0001 (two-way ANOVA with Tukey’s correction for multiple comparison. Statistical significance between mock and edited samples is depicted in the graph). **G.** CFC frequency for control and edited samples. Data are expressed as mean ± SEM (n = 2 biologically independent experiments, 2 donors) (two-way ANOVA with Dunnet correction for multiple comparisons; not significant). **H.** ddPCR analysis of *BCL11A* expression in erythroblasts derived from edited SCD HSPCs at day 13 of erythroid differentiation. *BCL11A* mRNA expression was normalized to *GAPDH* mRNA. Data are expressed as mean ± SEM (n = 3 biologically independent experiment, 3 donors) *P ≤ 0.05; **P ≤ 0.01 (one-way ANOVA with Dunnett correction for multiple comparison. Statistical significance between mock and edited samples is depicted in the graph). **I.** RT-qPCR analysis of γ (^G^γ + ^A^γ)- and β^S^-globin mRNA in control and edited SCD erythroblast at day 13 of erythroid differentiation. γ- and β^S^-globin mRNA expression was normalized to α-globin mRNA and expressed as a percentage of the γ- + β^S^-globin mRNA. The EE ± SEM is indicated for each sample in the lower part of the panel. Data are expressed as mean ± SEM (n = 3 biologically independent experiment, 3 donors). **P ≤ 0.01; ***P ≤ 0.001; ****P ≤ 0.0001 (two-way ANOVA with Dunnett correction for multiple comparisons. Statistical significance between mock and edited samples is depicted in the graph). **J.** Correlation between HbF expression and total editing efficiency in erythroid samples derived from edited SCD HSPCs (erythroblasts and pools of BFU-E; n = 3 biological independent experiments, 3 donors). HbF expression was measured by cation-exchange HPLC and calculated over the total Hb tetramers. The background level measured in the control was subtracted from all the conditions. Total editing efficiency was calculated by adding the base editing and Cas9 editing efficiency (determined by Sanger sequencing at the individual sites) to the frequency of the 3.2-kb deletion and inversion detected by ddPCR. Base-editing efficiency was calculated using EditR software and Cas9-editing efficiency (InDels) was calculated using TIDE in samples subjected to Sanger sequencing. Statistical significance are shown in the table *P ≤ 0.05; **P ≤ 0.01 (Multiple t test).

Overall, in erythroid liquid cultures and pools of BFU-E multiplex editing led to greater down-regulation of *BCL11A-XL* (encoding the isoform responsible for HbF silencing) and an increase of γ-globin production compared to single editing, regardless of the specific editor used (**Figure 4H-I** and **Supplementary Figure 3A-B and M**). To circumvent the disparity in editing efficiency between the samples, we correlated HbF reactivation and editing efficiency obtained in erythroid liquid cultures and pools of BFU-E (**Figure 4J** and **Supplementary Figure 4A-D**). This analysis confirmed that multiplex base editing correlated with stronger HbF reactivation compared to single editing. On the contrary, no additional effect was observed when targeting both BSs with the Cas9 nuclease compared to editing the +58-kb enhancer alone (**Supplementary Figure 4A-D**). A comparison of multiplex edited samples showed a significantly higher potency of DBE, followed by CBEs, ABEs, and lastly Cas9 (**Figure 4J**).

### CBE- and DBE-mediated multiplex targeting of BCL11A efficiently rescues the sickling phenotype

We selected CBE and DBE for further efficacy and safety analyses, given the higher HbF reactivation achieved with the +58 CBEI/+55 CBEII and +58 DBEI/+55 DBEI profiles compared to the ABE profiles. SCD HSPCs were co-transfected with the AncBE4max CBE-^30^ or the TadDE DBE-^31^encoding mRNAs and the same sgRNAs. Of note, while most of the enzymes used in the screening phase (**Figure 1 and 2 and Supplementary Figure 1**) are nearly PAM-less, both these base editors recognize a protospacer followed by an NGG PAM, to reduce the occurrence of off-targets ^32^. Control and edited HSPCs were differentiated towards the erythroid lineage. Flow cytometry analysis of enucleation and early and late erythroid markers, showed no differences between control and edited samples (**Supplementary Figure 5**).

Editing efficiency in erythroid liquid cultures was similar with CBE and DBE at both GATA1 and ATF4 BSs. In particular, in the +58-kb enhancer 64.6% ± 5.3 and 67.0% ± 9.6 of editing efficiency were obtained with CBE and DBE, respectively (**Figure 5A**). Regarding the +55-kb enhancer, an editing efficiency of 67.8% ± 5.4 and 58.4% ± 12.0 was observed for CBE and DBE, respectively (**Figure 5B**). NGS analysis confirmed high editing efficiency with the generation of predicted base modification for all the profiles (**Supplementary Figure 6A-D**). Rare C>G and InDel events were observed at similar frequency when generating the +58 CBEI/+55 CBEII and the +58 DBEI/+55 DBEI profiles (**Figure 5C-D; Supplementary Figure 6A-H**). As previously observed, a higher frequency of 3.2-kb deletion was observed when generating the +58 DBEI/+55 DBEI profile compared to +58 CBEI/+55 CBEII profile (1.2% ± 0.5 with DBE and 0.3% ± 0.2 with CBE, **Figure 5E)**.

**Figure 5.**
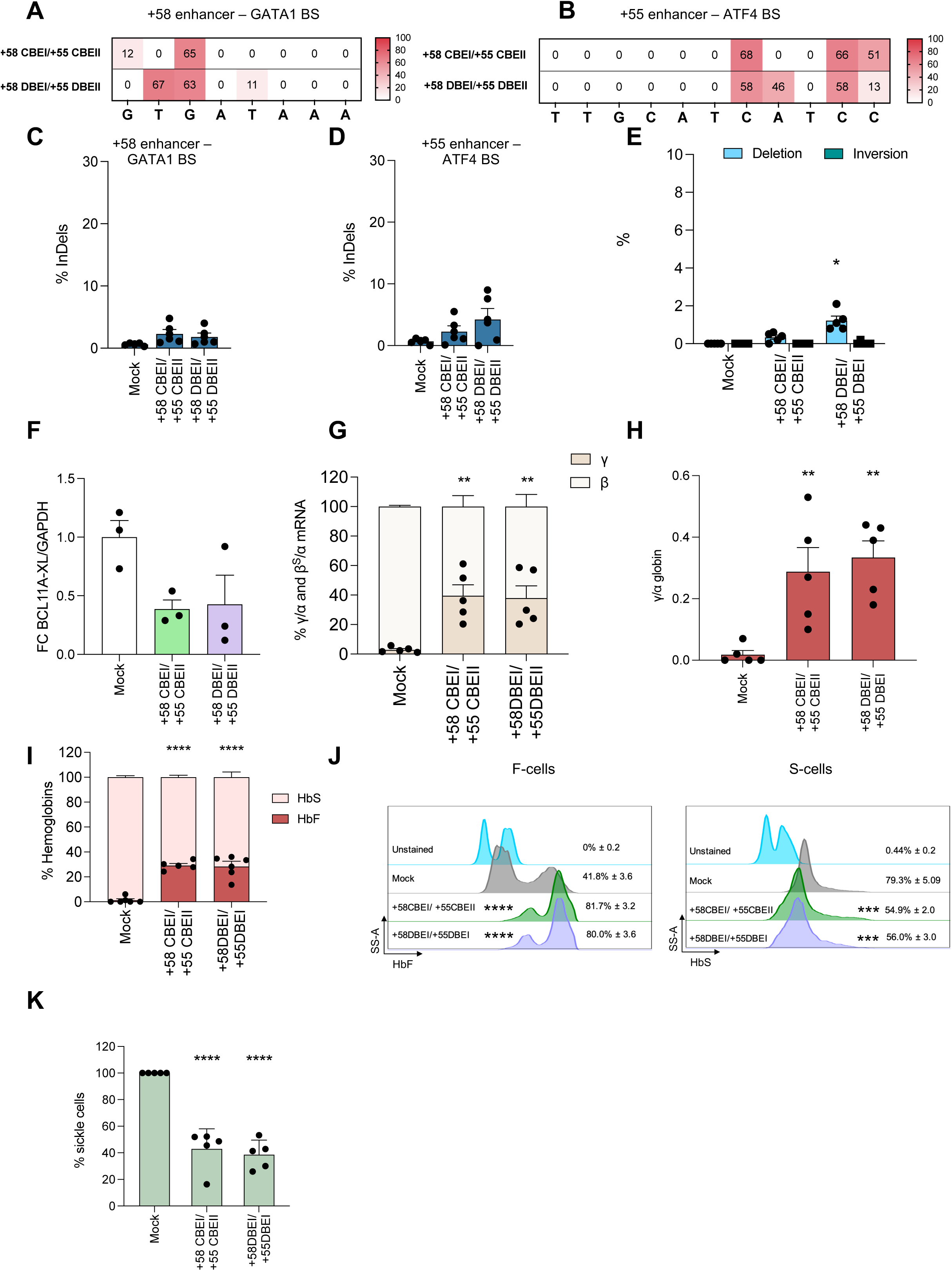
Multiplex CBE and DBE base editing in SCD HPSC-derived erythroblasts. **A-B**. C-G to T-A or/and A-T to G-C base-editing efficiency in the +58-kb (**A**) or +55-kb (**B**) regions calculated by EditR software in samples subjected to Sanger sequencing in erythroblasts derived from edited SCD HSPCs. GATA_bs_1 and ATF4_bs_2 were used in combination with AncBE4max to generate the +58 CBE/ +55 CBEII profile and in combination with TadDE to generate the +58 DBEI/ +55 DBEI profile. Data are expressed as mean (n = 1 biologically independent experiment, 5 donors). **C-D**. Frequency of InDels in the +58-kb (**C**) or +55-kb (**D**) regions measured by TIDE analysis, in samples subjected to Sanger sequencing in erythroblasts derived from edited SCD HSPCs. Data are expressed as mean ± SEM (n = 1 biologically independent experiment, 5 donors) (one-way ANOVA; not significant). **E.** Frequency of the 3.2-kb deletion/inversion, measured by ddPCR, in erythroblasts derived from edited SCD HSPCs. Data are expressed as mean ± SEM (n = 1 biologically independent experiment, 5 donors). *P ≤ 0.05 (one-way ANOVA with Dunnett correction for multiple comparison. Comparison of controls vs edited samples). **F.** ddPCR analysis of *BCL11A* mRNA in erythroblasts derived from edited SCD HSPCs at day 13 of *in vitro* erythroid differentiation. *BCL11A* mRNA expression was normalized to *GAPDH* mRNA. Data are expressed as mean ± SEM (n = 1 biologically independent experiment, 5 donors) (one-way ANOVA; not significant). **G.** RT-qPCR analysis of γ (^G^γ + ^A^γ)- and β^S^-globin mRNA in control and edited SCD patient’s erythroblasts at day 13 of erythroid differentiation. γ- and β^S^-globin mRNA expression was normalized to α-globin mRNA and expressed as a percentage of the γ- + β^S^-globin mRNA. Data are expressed as mean ± SEM (n = 1 biologically independent experiment, 5 donors). **P ≤ 0.01 (two-way ANOVA with Dunnett correction for multiple comparisons. Statistical significance between mock and edited samples is depicted in the graph). **H.** Expression of γ (^G^γ + ^A^γ)-globin chains measured by RP-HPLC in RBCs derived from SCD HSPCs. γ-globin expression was normalized to α-globin. Data are expressed as mean ± SEM (n=1 biologically independent experiment, 5 donors). **P ≤ 0.01 (one-way ANOVA with Dunnett correction for multiple comparisons. Statistical significance between mock and edited samples is depicted in the graph). **I.** Analysis of HbF and HbS by cation-exchange HPLC in RBCs derived from edited SCD HSPCs. We calculated the percentage of each Hb type over the total Hb tetramers. Data are expressed as mean ± SEM (n = 1 biologically independent experiment, 5 donors). ****P ≤ 0.0001 (two-way ANOVA with Dunnett correction for multiple comparisons. Comparison of mock vs edited samples). **J.** Representative flow cytometry histograms showing the percentage of HbF-expressing cells (F-cells) and HbS-expressing cells (S-cells) in CD235a^+^ population for unstained (CD235a stained only), control and edited samples. Data expressed as mean ± SEM is reported (n = 1 biologically independent experiment, 5 donors). ***P ≤ 0.001; ****P ≤ 0.0001 (two-way ANOVA with Dunnett correction for multiple comparisons. Statistical significance between mock and edited samples is depicted in the graph). **K.** Frequency of sickling cells upon O_2_ deprivation in mock and edited samples after normalization to the mock. Data are expressed mean ± SEM (n = 1 biologically independent experiment, 5 donors). ****P ≤ 0.0001 (one-way ANOVA with Dunnett correction for multiple comparisons. Statistical significance between mock and edited samples is depicted in the graph).

Following a marked reduction in *BCL11A-XL* mRNA observed in the edited samples compared to the controls (**Figure 5F**), a significant and consistent reactivation of γ-globin at the mRNA level was detected in the edited samples. Specifically, 39.5% ± 16.7% and 38.0% ± 18.4% of mRNAs encoding γ-globin were detected in HSPCs-derived cells carrying the +58 CBEI/+55 CBEII and +58 DBEI/+55 DBEI profiles, respectively (**Figure 5G**). These data were also confirmed at the protein level in mature RBCs by analyzing both globin chains and hemoglobin tetramers, with HbF accounting for 29.1% ± 3.6 and 28.3% ± 9.8 of total hemoglobin in the +58 CBEI/+55 CBEII and +58 DBEI/+55 DBEI profile, respectively (**Figure 5H-I**). Similarly, an increase of HbF-expressing cells was observed by generating the +58 CBEI/+55 CBEII and +58 DBEI/+55 DBEI profiles (**Figure 5J**), and importantly, a significant and homogeneous reduction in the frequency of HbS-expressing cells was observed in the edited samples compared to controls (**Figure 5J**). Finally, only 43.0% ± 15.0 and 38.7% ± 10.5 of sickle cells were detected in CBE- and DBE-edited samples, respectively (**Figure 5K**). These frequencies are similar to those observed in heterozygous, asymptomatic SCD carriers.

Overall, these data showed the efficacy of our strategy in reactivating HbF and rescuing the sickle phenotype.

### CBE- and DBE-mediated multiplex targeting of BCL11A: genotoxicity assessment

To evaluate the integrity of the locus, we performed long-read sequencing of the region encompassing the GATA1 and ATF4 BSs. We generated the +58/+55 Cas9 profile and compared it with our base editing profiles. Editing efficiencies between 27% and 43% were achieved in both BSs when generating the +58 CBEI/+55 CBEII, +58 DBEI/+55 DBEI and +58/+55 Cas9 profiles (**Figure 6A and B**). As expected, lower frequencies of alleles with 3.2-kb deletion/inversion were observed when using BEs compared to Cas9 nuclease (**Figure 6C**). For the remaining alleles, InDels associated with the generation of DSBs at the target sites were more frequent for Cas9 nuclease than BEs (**Figure 6C**). In addition, when using BEs, the majority of InDels were small (<50 bp) except for the +58 DBEI/+55 DBEI profile that showed 3.2% of large deletion (>200bp) at the GATA1 BS, (**Figure 6C**). Moreover, long-read sequencing showed a higher frequency of alleles simultaneously edited at GATA1 and ATF4 BSs when generating the +58 DBEI/+55 DBEI compared to the +58 CBEI/+55 CBEII profile (**Figure 6D**). To identify sgRNA-dependent potential off-target sites, we used *in silico* tools, and we also performed an unbiased genome wide analysis, GUIDE-Seq, in K562 cells. Interestingly, the top ten predicted off-target sites mapped to non-coding sequences and harbored more than 3 mismatches (**Figure 6E**). NGS of the top 5 GUIDE-seq predicted off-target and of the top 5 *in silico* predicted off-targets (**Supplementary Table 1**) in erythroblasts derived from edited SCD HSPCs validated few off-targets, which were limited to intronic and intergenic regions, suggesting no impact at protein level (**Figure 6F and G**). Additionally, off-target activity did not affect the expression of targeted genes in HSPCs (**Supplementary Table 2)**. Notably, although on-target activity was similar, CBE showed higher off-target activity compared to DBE, suggesting a higher specificity of the latter enzyme (**Figure 6E**). Importantly, no InDels were detected at the different off-target sites (**Figure 6F and G**).

**Figure 6.**
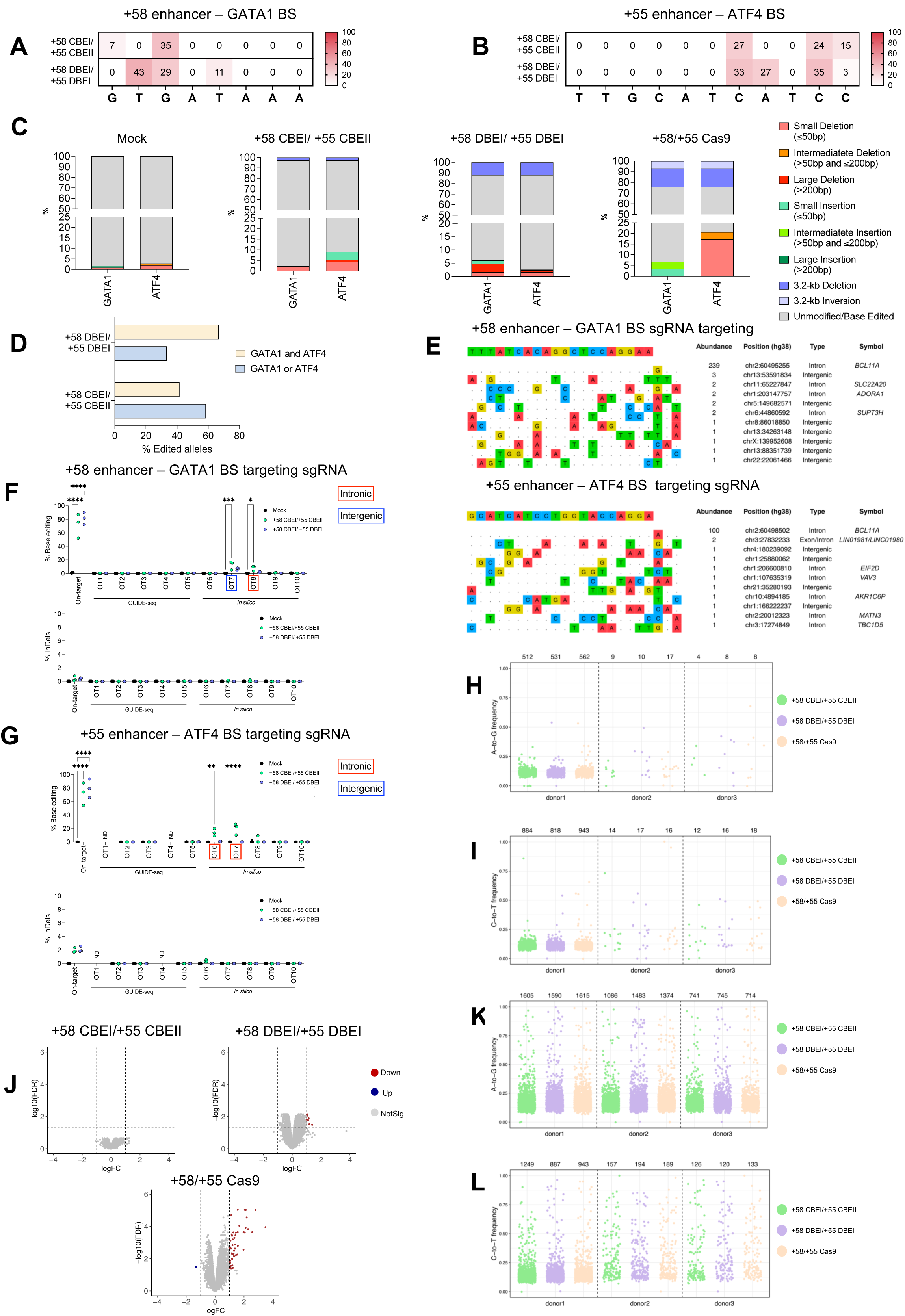
Evaluation of on-target aberrations and off-target activity of multiplex base editing of +58-kb and +55-kb regions. **A-B**. C-G to T-A or/and A-T to G-C base-editing efficiency in the +58-kb (**A**) and +55-kb (**B**) target region obtained with the generation of +58 CBEI/+55 CBEII profile, +58 DBEI/+55 DBEII and +58/+55 Cas9 profile. Editing efficiency in SCD HSPC-derived erythroblasts was evaluated by Sanger sequencing reaching 39% at GATA1 BS and ATF4 BS when generating the +58 CBEI/+55 CBEII profile and 50% and 38% at GATA1 and ATF4 BS, respectively when generating the +58 DBEI/ +55 DBEI profile. 25% and 13% of InDels were obtained at GATA1 and ATF4 BS, respectively, when using Cas9 nuclease. **C**. Comprehensive allele frequencies of diverse types of genomic rearrangements and InDel types in SCD HSPC-derived erythroblasts. We reported frequencies of events observed at individual GATA1 and ATF4 BSs (in the absence of the 3.2-kb deletion/inversion) and the frequencies of 3.2-kb deletion/inversion (GATA1-ATF4). InDels were classified as small, intermediate and large according to their size ([4-50bp], [51-200bp], >200bp). Del, deletion; Ins, insertion; Inv, inversion. Unmodified, no deletion, inversion or insertion observed **D.** Frequency of alleles edited at the GATA1 and ATF4 BSs, either simultaneously (GATA1 and ATF4) or at only one of the sites (GATA1 or ATF4). The reported data were adjusted for background error typical of this analysis by subtracting the frequency of edited alleles in the mock condition from the fraction of edited alleles in the edited conditions. **E**. sgRNA-dependent off-target sites of the GATA_bs_1 (top panel) and ATF4_bs _2 (low panel) sgRNAs in K562 cells, as evaluated by GUIDE-seq analysis. sgRNAs were coupled with a Cas9-nuclease corresponding to the Cas9 nickase included in base editor (Cas9-NGG). The protospacer targeted by each sgRNA is reported on top of each panel, followed by the off-target sites and their mismatches with the on-target (highlighted in color). The number of sequencing reads (abundance), the chromosomal coordinates (hg38) the type of mapped site and the symbol of the gene, of each off-target site are reported. The cutoff was set to a level of detection of at least three reads. **F-G**. Base editing and InDel frequency at on-target and off-target (OT) sites, in SCD HSPC-derived erythroblasts, evaluated for GATA1 BS targeting sgRNA (**F**) and ATF4 BS targeting sgRNA (**G**) for control, +58 CBEI/+ 55 CBEII and +58 DBEI/+55 DBEI profiles as measured by targeted NGS. OT sites in introns or intergenic regions for which we detected editing are highlighted by a red or blue squared shape, respectively. Data are expressed as individual values and median (n=3 biologically independent experiments, 3 donors) *P≤0.05, **P ≤ 0.01, ***P ≤ 0.001 and ****P≤0.0001 (two-way ANOVA with Dunnet’s Statistical significance between mock and edited samples is depicted in the graph). **H-I.** Strip plots showing the variant allele frequency (A > G, **H**; C > T, **I**) in exons, observed in SCD HSPCs obtained from three different donors and measured by WES. The total number of variants are indicated above each sample. **J**. Volcano plots showing differential gene expression between cells treated with CBE, DBE, or Cas9 nuclease and mock-treated samples. RNA-seq was performed 48 h after electroporation in SCD HSPCs from 3 different donors. The horizontal dashed line indicates the threshold on the false discovery rate (FDR ≤ 0.05), and the vertical dashed lines correspond to the threshold on log2FC ≥ 1 or ≤ -1. Upregulated genes are indicated in red, and downregulated genes are in blue. Genes in grey are not differentially expressed. **K-L.** Strip plots showing the variant allele frequency (A > G, **K**; C > T, **L**) in the transcriptome observed in SCD HSPCs obtained from three different donors and measured by RNA-seq. The total number of variants are indicated above each sample.

We also performed whole-exome sequencing (WES) analysis in samples harboring the +58 CBEI/+55 CBEII, +58 DBEI/+55 DBEI and +58/+55 Cas9 profiles. In general, a similar frequency of A-to-G and C-to-T mutations was detected in the different samples, indicating that base editors do not increase the mutational burden (**Figure 6H and I**).

To examine the effect of base editing on the overall gene expression profile, we performed RNA-seq of SCD HSPCs mock-edited or edited with DBE, CBE, or Cas9 nucleases targeting both +58-kb and +55-kb enhancers. Overall, we observed few or no differentially expressed genes (DEGs) in DBE- and CBE-treated samples, respectively (**Figure 6J and Supplementary Table 2,** log2 fold change (log2FC) ≥ 1 or ≤-1; false discovery rate (FDR) ≤ 0.05). On the contrary, 49 genes were upregulated in Cas9-treated samples (**Figure 6J**), mainly involved in p53 pathways, inflammatory response, and apoptosis. Finally, the frequency of A-to-G and C-to-T conversions in the transcriptome was comparable across all edited samples, indicating that base editors do not increase RNA deamination (**Figure 6K and L**).

Overall, these data showed that CBE- and DBE-mediated multiplex targeting of *BCL11A* is potentially a safer approach in terms of genotoxicity compared to the CRISPR/Cas9 nuclease treatment.

### Efficient multiplex base editing of *BCL11A* enhancers in repopulating HSCs

To evaluate the ability of BEs to simultaneously target the *BCL11A* enhancers in repopulating HSCs, we xenotransplanted control and CBE- or DBE-edited HD HSPCs into immunodeficient NBSGW mice (**Figure 7A**). Sixteen to seventeen weeks post-transplantation, no differences were observed between edited and control HSPCs with regard to engraftment and differentiation potential, as measured by the frequency of human CD45^+^ cells in hematopoietic tissues and the proportion of the different lineages (**Figure 7B and Supplementary Figure 7A**). The base editing efficiency in human BM cells (33.7% ± 4.6/59.2% ± 13.3 for +58 CBEI/+55 CBEII, and 68.8% ± 15/63.1% ± 9.4 for +58 DBEI/+55 DBEII) was similar to the frequency observed in the input cells and in the other hematopoietic compartments (blood, erythroid precursors, thymus, spleen and B-cells; **Figure 7C-F**). However, CBE editing efficiency at the +58 enhancer tends to be reduced *in vivo* compared to the input HSPCs (**Figure 7C**). InDels and 3.2-kb deletion were detected at low frequencies in engrafted cells (**Supplementary Figure 7B and C**). Notably, a significant reduction of 3.2-kb deletion frequency was detected *in vivo* compared to the input cells (**Supplementary Figure 7C**).

**Figure 7.**
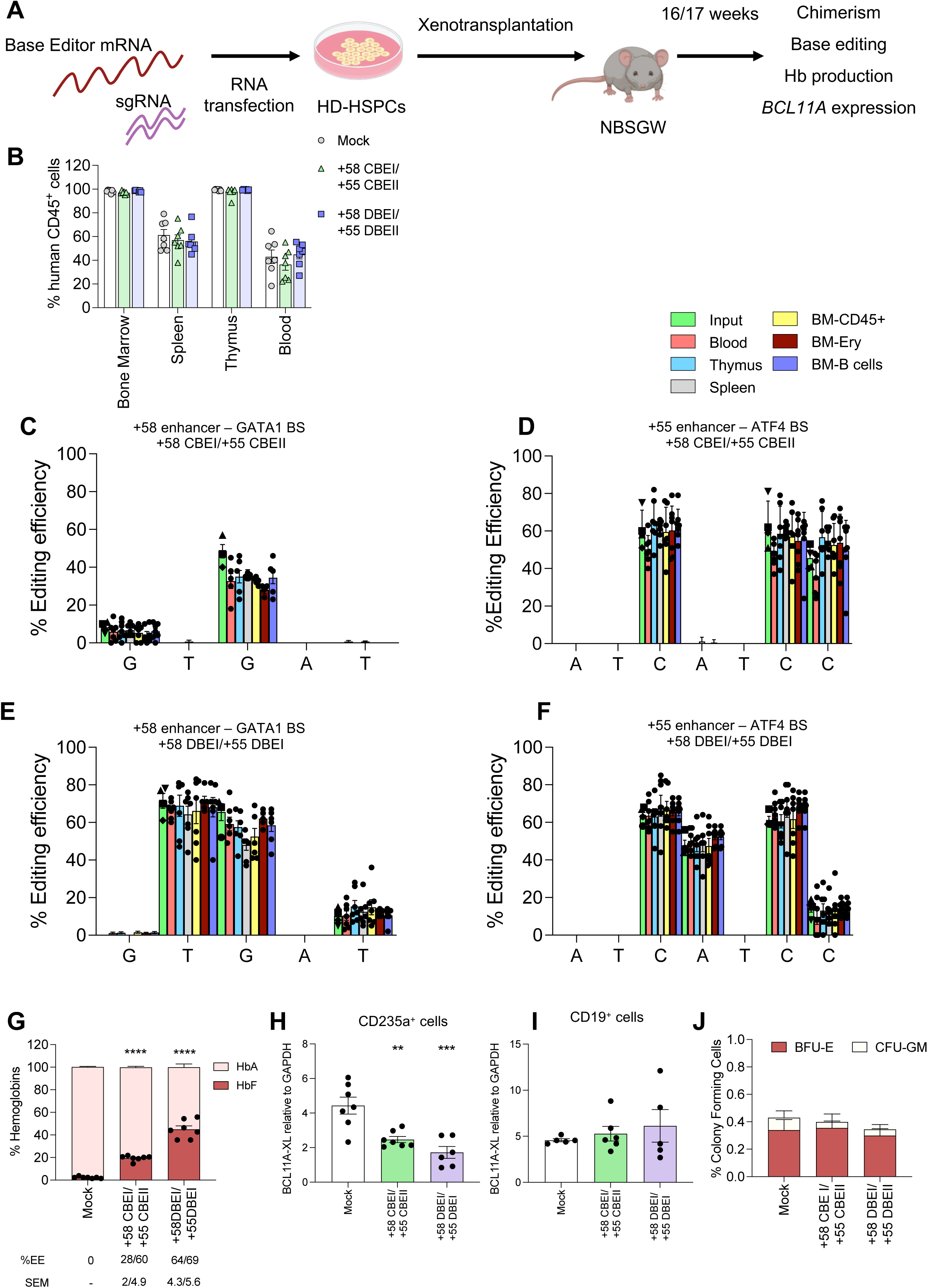
Simultaneous base editing of *BCL11A* enhancers in repopulating HSCs. **A**. Experimental protocol used for HD HSPC xenotransplantation experiments. BE mRNAs and sgRNAs were co-transfected in HD HSPCs, and control and edited cells were xenotransplanted into NBSGW immunodeficient mice 1 day after transfection. Mice were euthanized 16/17 weeks after transplantation and their hematopoietic tissues and organs were collected and analyzed. **B**. Engraftment of human cells in NBSGW mice transplanted with control (mock-transfected) or edited HSPCs 16/17 weeks post-transplantation (n = 7 mice per group). Engraftment is represented as the percentage of human CD45^+^ cells in the total murine and human CD45^+^ cell population in BM, spleen, thymus, and peripheral blood. Each data point represents an individual mouse. Data are expressed as mean ± SEM. (two-way ANOVA; not significant). **C-F**. C-G to T-A or/and A-T to G-C base-editing efficiency, calculated using EditR for +58 CBEI/ +55 CBEII (**C** and **D**) and +58 DBEI/ +55 DBEI (**E** and **F**) profiles at the +58-kb (**C** and **E**) and +55-kb (**D** and **F**) regions in the input, blood-, bone marrow (BM)-, thymus-, spleen-derived human samples subjected to Sanger sequencing. BM samples include CD45^+^ cells, B cells and erythroid cells (Ery). Each data point represents an individual mouse. The frequency of base editing in the input was calculated in cells cultured in the HSPC medium (▪), in liquid erythroid cultures (▴), and in pools of BFU-E (◆) and CFU-GM (▾). Data are expressed as mean ± SEM [n = 1 biologically independent experiment (Input), n = 7 mice per group]. **G**. Analysis of HbF and HbS by cation-exchange HPLC in human CD235a^+^ BM-sorted erythroid cells from mice transplanted with control and edited HSPCs, 16/17 weeks post-transplantation. We calculated the percentage of each Hb type over the total Hb tetramers. Data are expressed as mean ± SEM (n = 7 mice per group). Each data point represents an individual mouse. ****P ≤ 0.0001 (two-way ANOVA with Dunnett correction for multiple comparisons. Statistical significance between mock and edited samples is depicted in the graph). **H-I**. RT-qPCR analysis of *BCL11A* in BM-sorted CD235a^+^ (**I**) or CD19^+^ (**J**) cells from mice transplanted with control and edited HSPCs, 16/17 weeks post-transplantation. *BCL11A* mRNA expression was normalized to *GAPDH* mRNA. Data are expressed as mean ± SEM (n = 5 to 7 mice per group). Each data point represents an individual mouse. **P ≤ 0.01; ***P ≤ 0.001 (One-way ANOVA with Tukey’s correction for multiple comparisons. Statistical significance between mock and edited samples is depicted in the graph). **K.** Human hematopoietic progenitor content in BM human CD45^+^ cells derived from mice transplanted with control and edited HSPCs. We plotted the percentage of human CD45^+^ cells giving rise to BFU-E and CFU-GM. Data are expressed as mean ± SEM (n =mice per group). (two-way ANOVA; not significant).

Human CD235a^+^ erythroid cells sorted from BM of mice transplanted with edited cells, showed increased HbF expression at the RNA and protein levels compared to control groups (**Figure 7G, and Supplementary Figure 7D and E**). Notably, higher HbF reactivation was observed in samples carrying the +58 DBEI/+55 DBEI profile, which showed the highest base editing efficiency (**Figure 7G, and Supplementary Figure 7D and E**). Correlation analysis between HbF expression and editing efficiency in BM CD235a^+^ sorted cells also confirmed a more potent HbF reactivation associated with the +58 DBEI/+55 DBEII profile *in vivo* compared to the +58 CBEI/+55 CBEII profile (**Supplementary Figure 7F**). Importantly, *BCL11A* down-regulation was observed in edited human BM CD235a^+^ erythroid cells (**Figure 7H**), while no differences were detected between control and edited samples in BM human non-erythroid CD19^+^ B-cells (**Figure 7I**).

Human CD45^+^ BM cells were isolated and subjected to a CFC assay. Control and edited samples showed a similar number of erythroid and granulo-monocytic colonies, demonstrating no impact of base editing on the clonogenic potential of engrafted human cells (**Figure 7J**). Editing efficiency in BFU-E and CFU-GM was similar to that measured in the input populations except for the GATA1 BS in the +58-kb enhancer in CBE-treated samples, as observed in human CD45^+^ BM cells (**Supplementary Figure 7G-J**). Furthermore, InDels and 3.2-kb deletions were detected at low frequencies (**Supplementary Figure 7K and L**). *BCL11A-XL* downregulation was observed in BFU-E (**Supplementary Figure 7M**), resulting also in globin reactivation at RNA and protein levels (**Supplementary Figure 7N-P**).

Overall, these results suggest high efficiency and a good safety profile of our multiplex base editing approach in HSCs and their differentiated progeny.

## Discussion

Since the discovery of elements controlling the expression of HbF, a variety of genome editing strategies based on the disruption of cis- or trans-regulatory elements of *HBG1/2* to induce HbF reactivation have been developed for the treatment of β-hemoglobinopathies ^7,8,11,14^.

Importantly, Cas9 nuclease-mediated disruption of the +58-kb erythroid-specific *BCL11A* enhancer has been recently approved as a clinical therapy for patients with SCD and β-thalassemia ^9,10^. However, although significant HbF reactivation was achieved, the clinical study showed variability between the patients in HbF levels. Furthermore, SCD patients retained high HbS levels and showed a modest correction of ineffective erythropoiesis. Thus, further optimization of the strategy is advisable to reach the complete rescue of the phenotype by reaching higher levels of HbF. In addition, the generation of Cas9 nuclease-induced DSBs or large genomic rearrangements can be detrimental to HSCs ^12,33–36^. Therefore, it has been lately proposed to use the base editing technology to disrupt the GATA1 activator BS in the erythroid-specific *BCL11A* enhancer without creating InDels, and reactivate HbF in the progeny of SCD and β-thalassemia patient HSPCs ^37–39^.

Here, we used SCD HSPCs to perform a screening of different BE/sgRNAs to disrupt the GATA1 and ATF4 activator BS in the erythroid-specific *BCL11A* enhancer regions (+58-kb and +55-kb, respectively) by changing specific targeted nucleotides. Indeed, modifying the GATA1 BS differentially reduces the ability of GATA1 to bind to the *BCL11A* enhancer, and thus modulate gene expression ^29^. Importantly, we were able to create different editing profiles for both motifs, by combining different sgRNAs and ABEs or CBEs to generate A>G or C>T base conversions. Additionally, the use of TadDE allowed us to extend the number of edited bases and further disrupt the GATA1 and ATF4 BSs.

When targeting individually the GATA1 and ATF4 BSs, only the generation of the +58 CBEI profile at the GATA1 BS led to HbF expression and consequent rescue of the sickle phenotype, similar to the approved Cas9 nuclease-based approach. Thus, to further increase HbF levels, we hypothesized that combined editing of the +58-kb and +55-kb enhancers could lead to a stronger reduction of *BCL11A* expression, and consequently greater HbF induction compared to editing a single enhancer alone. Of note, multiplex targeting of the +58-kb and +55-kb *BCL11A* enhancers using CRISPR/Cas9 (although targeting different binding site locus in the +55-kb enhancer) was recently reported to efficiently reactivate HbF expression ^40,41^. However, the use of CRISPR/Cas9 nucleases cause potentially dangerous DSBs and large genomic rearrangements.

Correlation analysis on single cell-derived colonies allowed us to identify the critical bases involved in the binding of GATA1 and ATF4 at the *BCL11A* enhancers and the most potent profiles in terms of HbF reactivation that we can combine to further improve the pathological phenotype.

Regarding the +58-kb region, our findings corroborate previous studies that reported an important role of the G_3_ of the GATA binding motif in regulating GATA1 binding ^29^. This is supported by the elevated γ-globin levels observed upon generating the +58 CBEI profile (<u>G</u>T<u>G</u>ATAAA), effectively replicating the outcomes achieved with the approved Cas9 nuclease strategy. In addition, the A_6_ position in the GATA1 BS (GTGAT<u>A</u>AA; +58 ABEIII profile) plays an important role in GATA1 binding, as already observed in other works^29^ and confirmed by the strong increase of γ-globin re-activation. In contrast, targeting the positions A_6_ to A_8_ in the GATA1 binding motif (GTGAT<u>AAA</u>) to generate the +58 ABEII profile resulted in reduced γ-globin expression compared with the +58 ABEIII profile. This reduction suggests that the A>G conversion at the positions A_7_ and A_8_ in addition to the A_6_ base conversion might restore a binding motif for transcriptional activators, which affects *BCL11A* and, therefore, γ-globin expression

Similarly, T>C conversions generated at position T_2_ and T_5_ (G<u>T</u>GA<u>T</u>AAA) to create the +58 ABEI profile led to less potent γ-globin re-activation and presumably less efficient disruption of GATA1 BS compared to the generation of the +58 ABEIII profile, although previous studies reported high GATA1 displacement when T>C mutation was occurring at position 5 of the GATA binding motif compared to the targeting of A_6_ ^29^. These results suggest that site-specific requirements for GATA1 binding at the +58-kb enhancer, likely because the surrounding sequence, could also influence GATA1 occupancy .

Regarding the +55-kb region, the generation of the +55 CBEI profile led to C>T conversion of all the Cs (C_4,_ C_7,_ C_10_ and C_11_) of the ATF4 binding motif (TTG<u>C</u>AT<u>C</u>AT<u>CC</u>), while with the generation of +55 CBEII profile only C_7,_ C_10_ and C_11_ were converted (TTGCAT<u>C</u>AT<u>CC</u>). Our data suggest that the insertion of two points mutations at C_4_ and C_7_ (+55 CBEI profile) does not further displace ATF4 as compared to the generation of C_7_ mutation alone (+55 CBEII profile), indicating that the primary factor affecting ATF4 binding (and subsequently *BCL11A* expression and γ-globin reactivation), is the C>T conversion at C_7_. Conversions of the A_5_ and A_8_ of the ATF4 binding motif (TTGC<u>A</u>TCATCC, +55 ABEI and TTGC<u>A</u>TC<u>A</u>TCC, +55 ABEII) led to low *HBG* levels, suggesting a modest impact on ATF4 binding. However, the generation of the +55 ABEII profile (TTGC<u>A</u>TC<u>A</u>TCC) showed a higher effect in terms of γ-globin reactivation compared to the +55 ABEI profile (TTGC<u>A</u>TCATCC). This result can either indicate a stronger impact of A_8_ mutation in the displacement of ATF4 or that the simultaneous generation of A_5_ and A_8_ mutation enhanced the disruption of the ATF4 binding motif.

Finally, we used DBE to further disrupt the GATA1 and ATF4 BSs; however, *in vitro* we observed only a modest increase in HbF reactivation compared to the use of CBE alone, suggesting a minor role of As in the binding of these factors to the *BCL11A* enhancers.

Multiplex targeting of the +58-kb and +55-kb regions led to a significant reduction of *BCL11A* expression and consequently to a potent HbF reactivation. Importantly, HbF reactivation obtained upon multiplex DBE- and CBE-mediated base editing reached the levels needed to ameliorate the clinical manifestations of SCD, which have been defined as 70% of HbF-expressing cells and HbF accounting for 30% of the total hemoglobin for most of the patient samples^42^.

Importantly, fewer genomic rearrangements were detected in base-edited samples compared to cells treated with Cas9 nuclease, highlighting the better safety profile of our approach. We showed that large genomic rearrangements (particularly deletions) within the 3.2-kb region between the GATA1 and ATF4 occur with more frequently with DBE compared to the CBE, while ABE showed no such large deletions. While higher rearrangements with DBE can be explained simply by the additive effect of CBE and ABE activity, the reduced deletion frequency observed with ABE compared to CBE is probably due to an inefficient excision of inosines (in the case of ABEs) compared to the excision of the uracil base (in the case of CBEs), which reduces the generation of DSBs. In addition, the C>G base conversions that was observed with CBE and DBE have been previously associated with DSB generation, deletions and other large genomic rearrangements^43^. As for the 3.2-kb deletion, also on-target unintended large deletions (> 200 bp) were more frequent in DBE-treated samples (+58 DBEI/+55 DBEI profile) compared to CBE-treated samples (+58 CBEI/ +55 CBEII profile); however, deletion frequency remained lower compared to the samples treated with the Cas9 nuclease.

Targeted deep sequencing of the top 5 GUIDE-seq and *in* silico predicted off-targets in base-edited HSPCs showed a low sgRNA-dependent off-target activity, occurring in non-exonic regions, with no predicted effect at the protein level, and no consequences at RNA level, as confirmed by RNA-seq. In addition, a comparison between DBE and CBE revealed a higher specificity of DBE. Importantly, no indels were detected at the off-target sites, suggesting that base editors minimize the possibility of generating DSB-induced genomic rearrangements such as translocations. In addition, we comprehensively assessed the sgRNA-independent off-target DNA activity by WES, demonstrating no sgRNA-independent off-target activity within exons in DBE-and CBE-treated samples. Finally, we analyzed sgRNA-independent off-target RNA activity showing that proposed base editing strategies did not lead to deamination of the cellular transcriptome.

Xenotransplantation experiments of multiplex base-edited HSPCs in immunodeficient mice confirmed high editing efficiencies in long-term repopulating HSCs and their progeny, demonstrating that multiplex base editing does not affect engraftment and multilineage differentiation of HSCs. Of note, we achieved higher editing frequency and HbF reactivation with DBE compared to CBE *in vivo*. Correlation analysis between editing efficiency and HbF levels suggested that DBE-induced multiple mutations in the *BCL11A* enhancers better evict GATA1 and ATF4 and down-regulate *BCL11A in vivo*. These results also suggest that CBEs might be less efficient in *bona fide* HSCs. In addition, CBEs might have induced some toxicity *in vivo* which could have affect cell viability and self-renewal capacity of HSCs, as previously proposed ^14^.

Of note, complete inactivation of *BCL11A,* adversely affects lymphoid development and vital HSC functions ^44–46^. However, *in vivo* experiments confirmed previous observations that the downregulation of *BCL11A* achieved by targeting the GATA1 BS in the +58-kb erythroid-specific enhancer does not impair *BCL11A* expression in other lineages ^47^ . In addition, here we demonstrated that also the targeting of ATF4 BS in the +55-kb enhancer led to specific downregulation of *BCL11A* in the erythroid lineage without impacting its expression in B cells. Nonetheless, complete knock out of *BCL11A* can also harm the erythroid lineage by affecting the human RBC enucleation^48^, but no alteration in the enucleation rate were observed with our base editing strategy, likely because of the partial down-regulation of *BCL11A* expression.

In conclusion, this study provides insights into the applicability of multiplexed base editing strategies to treat SCD and potentially β-thalassemia patients through disruption of the erythroid-specific *BCL11A* enhancers in the +58-kb and +55-kb regions. It is worth noting that our approaches can serve as universal therapeutic strategies for both SCD and β-thalassemia patients, as they do not require the design of mutation-specific CRISPR-Cas9-based tools, as proposed in previous studies ^37,49,50^. The levels of HbF achieved with multiplex base editing were sufficient to ameliorate the sickling phenotype. The clinical translation of our approach will require a deeper assessment of the off-target activity of our base editing systems to ensure the safety of our approaches, the establishment of a large-scale transfection protocol with clinical-grade reagents as well as biodistribution studies in mice.

## Supplementary Figure legends

**Supplementary Figure 1.**
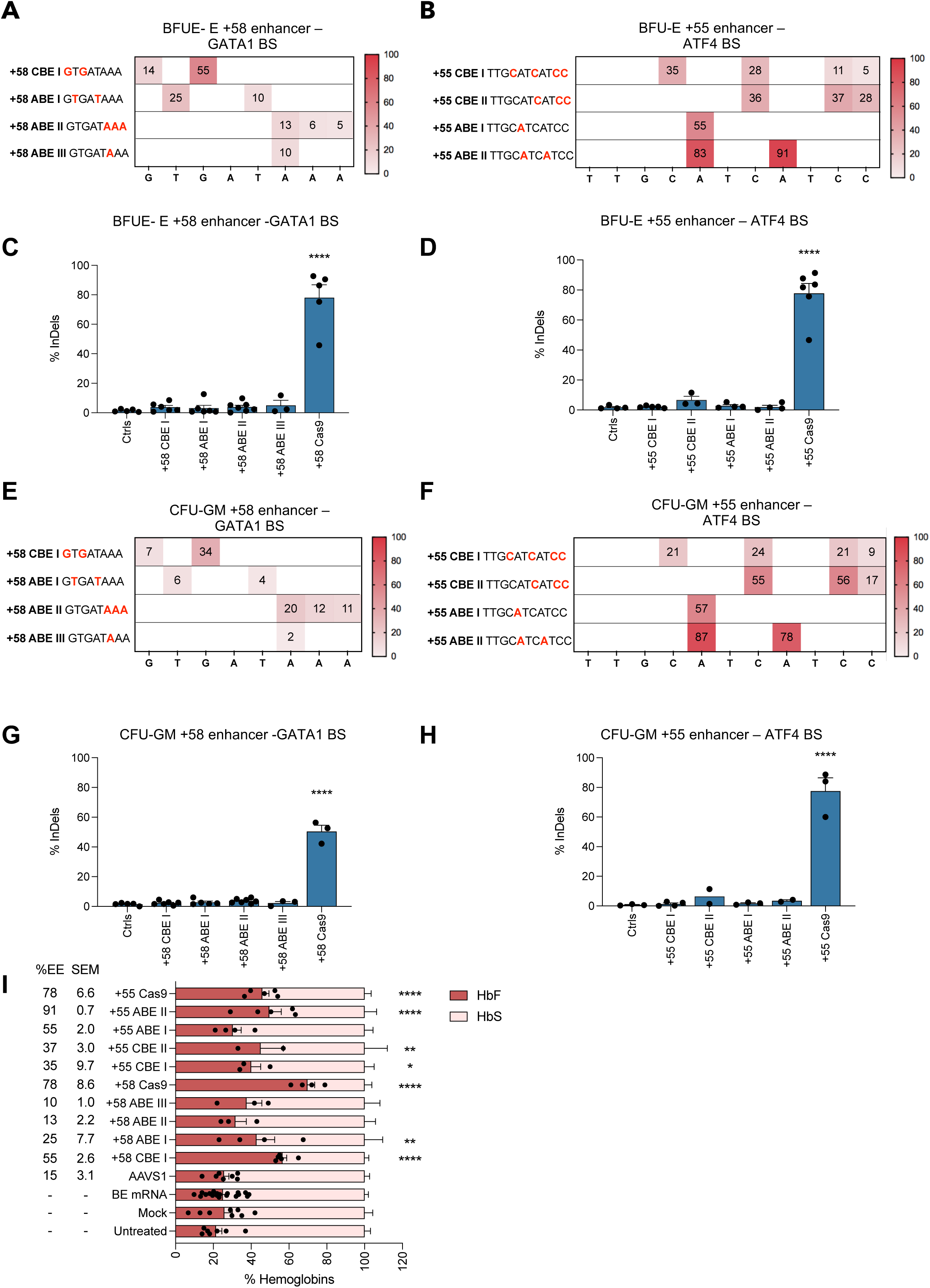
Base editing of the erythroid-specific *BCL11A* enhancers in SCD HPSC-derived BFU-E and CFU-GM colonies. **A-B**. C-G to T-A or A-T to G-C base-editing efficiency calculated by the EditR software in samples subjected to Sanger sequencing in pools of BFU-E colonies derived from SCD HSPCs edited in the +58-kb (**A**) or +55-kb (**B**) regions. Data are expressed as mean (n = 2 to 4 biologically independent experiments, 2 to 5 donors). **C-D**. Frequency of InDels, measured by TIDE analysis, in samples subjected to Sanger sequencing in pools of BFU-E colonies derived from SCD HSPCs edited in the +58-kb (**C**) or +55-kb (**D**) regions. Data are expressed as mean ± SEM (n = 2 to 4 biologically independent experiments, 2 to 5 donors). ****P ≤ 0.0001 (One-way ANOVA. Comparison of controls vs edited samples) **E-F**. C-G to T-A or A-T to G-C base-editing efficiency calculated by the EditR software in samples subjected to Sanger sequencing in pools of CFU-GM colonies derived from SCD HSPCs edited in the +58-kb (**E**) or +55-kb (**F**) regions. Data are expressed as mean (n = 2 to 4 biologically independent experiments, 2 to 5 donors). **G-H**. Frequency of InDels, measured by TIDE analysis, in samples subjected to Sanger sequencing in pools of CFU-GM colonies derived from SCD HSPCs edited in the +58-kb (**G**) or +55-kb (**H**) regions. Data are expressed as mean ± SEM (n = 2 to 4 biologically independent experiments, 2 to 5 donors). ****P ≤ 0.0001 (One-way ANOVA. Comparison of controls vs edited samples) **I**. Analysis of HbF and HbS by cation-exchange HPLC in pools of BFU-E colonies. We calculated the percentage of each Hb type over the total Hb tetramers. Data are expressed as mean ± SEM (n = 3 to 4 biologically independent experiments, 3 to 4 donors). *P ≤ 0.05; **P ≤ 0.01; ****P ≤ 0.0001 (two-way ANOVA with Dunnett correction for multiple comparisons. Statistical significance between mock and edited samples is depicted in the graph).

**Supplementary Figure 2.**
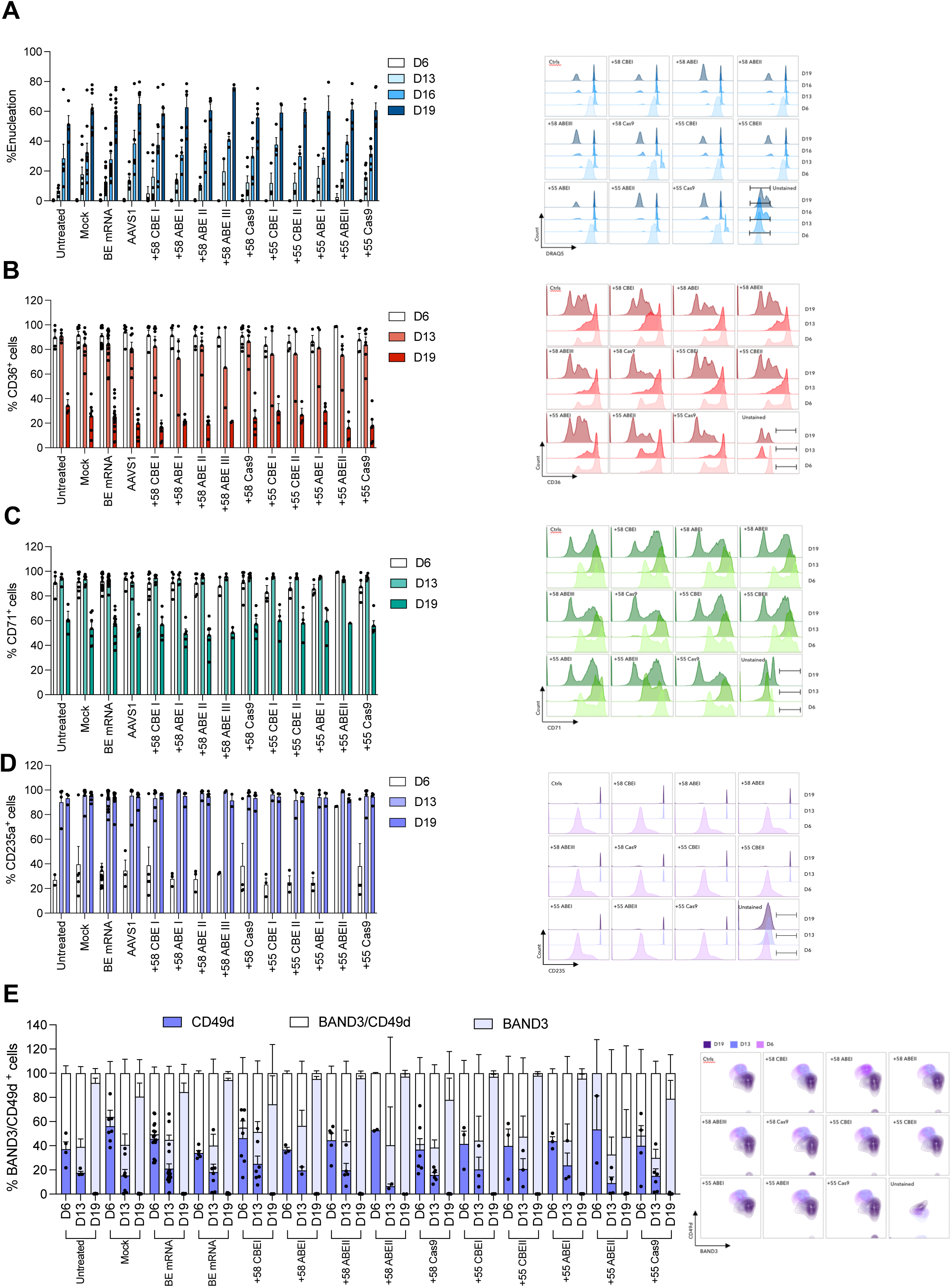
Erythroid differentiation of SCD HSPCs upon base editing of the erythroid-specific *BCL11A* enhancers. **A**. Frequency of enucleated cells at day 6, 13, 16 and 19 of erythroid differentiation, as measured by flow cytometry analysis of DRAQ5 nuclear staining in control and edited samples. Data are expressed as mean ± SEM (n = 2 to 6 biologically independent experiments, 2 to 5 donors). Representative flow cytometry histograms showing the DRAQ5^−^ cell population for control and edited samples are reported. **B-D**. Frequency of CD36^+^ (**B**), CD71^+^ (**C**) and CD235a^+^ (**D**) cells at day 6, 13 and 19 of erythroid differentiation, as measured by flow cytometry analysis of CD36, CD71 and CD235a erythroid markers. Data are expressed as mean ± SEM (n = 2 to 6 biologically independent experiments, 2 to 5 donors). Representative flow cytometry histograms showing the CD36^+^ (**B**), CD71^+^ (**C**) and CD235a^+^ (**D**) cell population for control and edited samples are reported. **E**. Frequency of CD49d^+^, BAND3^+^ and CD49d^+^/BAND3^+^ in 7AAD^−^/CD235a^+^ cells at day 6, 13 and 19 of erythroid differentiation, as measured by flow cytometry analysis of CD49d and BAND3 erythroid markers. Data are expressed as mean ± SEM (n = 2 to 6 biologically independent experiments, 2 to 5 donors). Representative flow cytometry contour plots showing the CD49d^+^, BAND3^+^ and CD49d^+^/BAND3^+^ cell population for control and edited samples are reported.

**Supplementary Figure 3.**
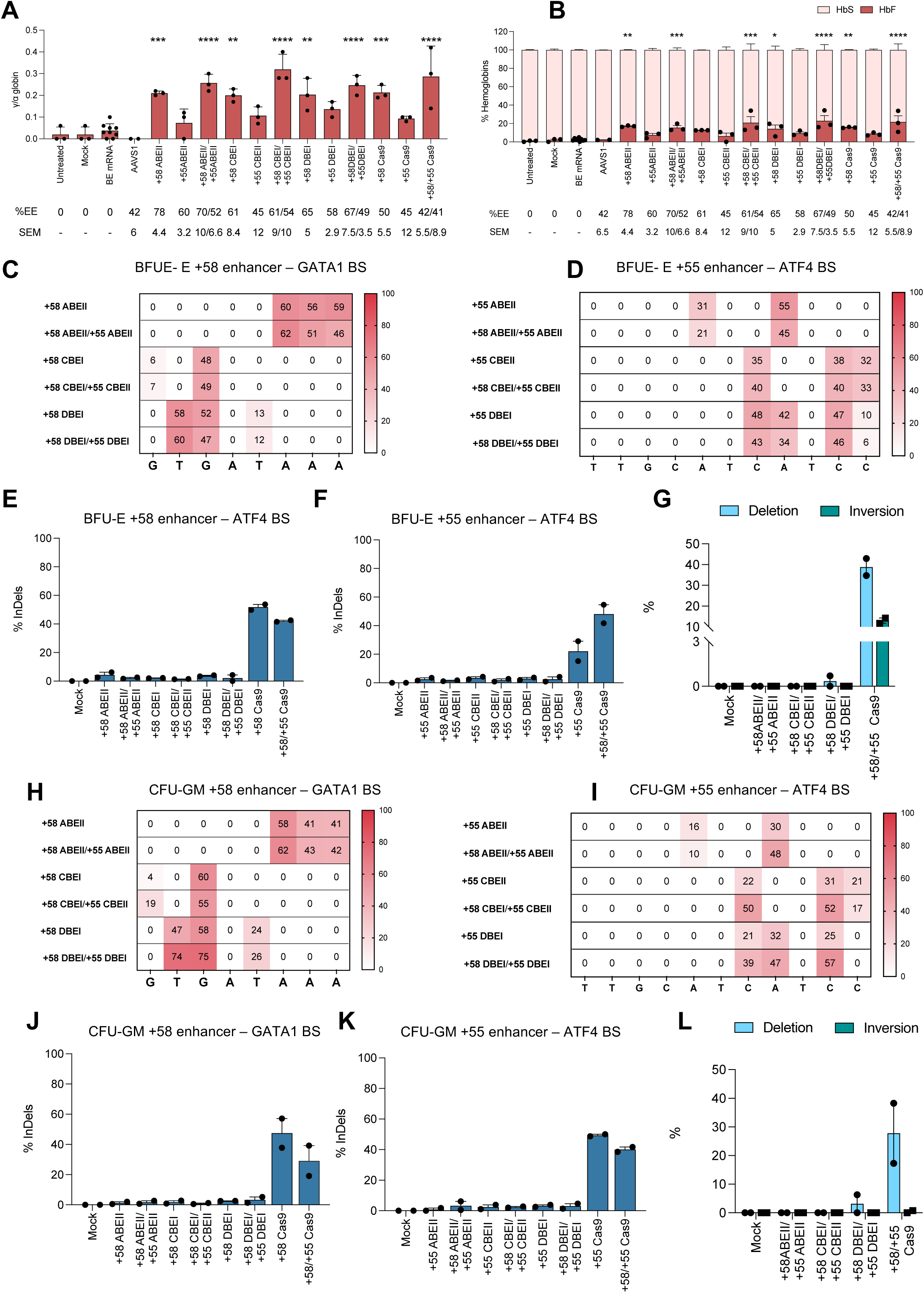
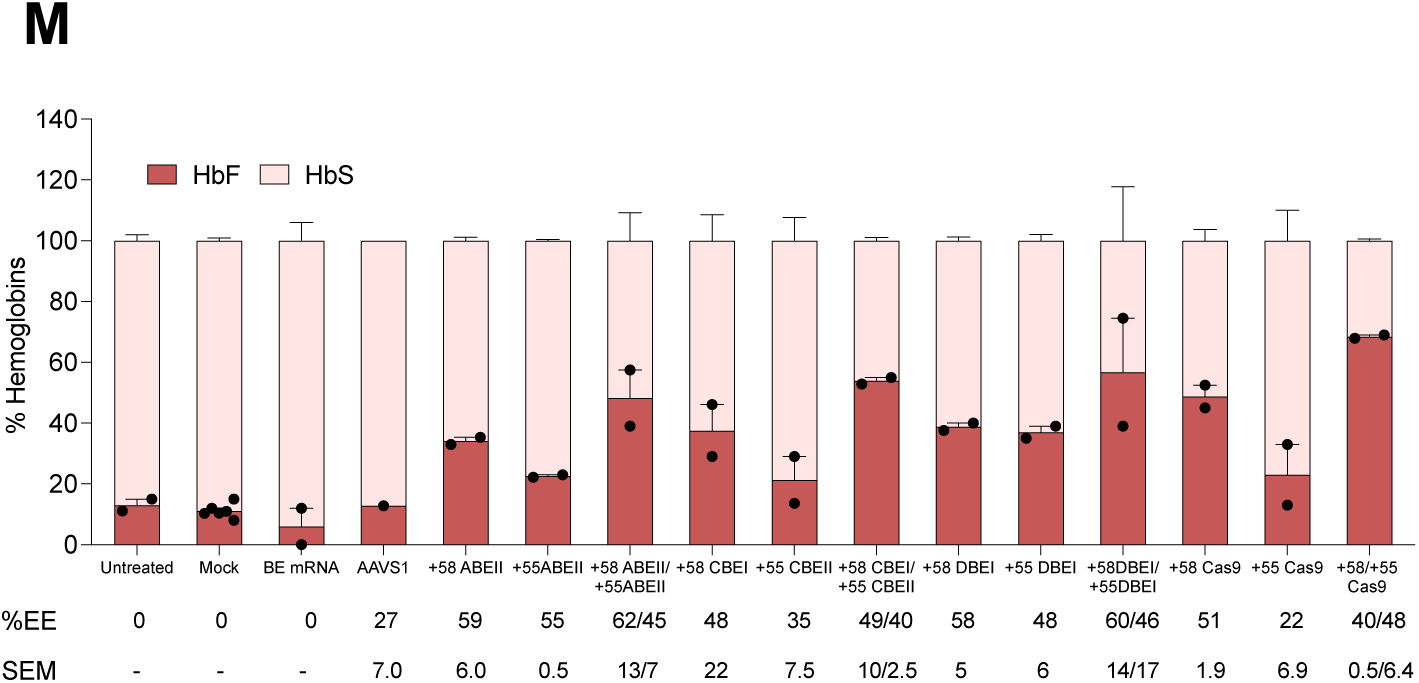
Multiplex base editing of the erythroid-specific *BCL11A* enhancers. **A.** Expression of γ (^G^γ + ^A^γ)-globin chains measured by RP-HPLC in RBCs derived from SCD HSPCs. γ-globin expression was normalized to α-globin. The EE ± SEM is indicated for each sample in the lower part of the panel. Data are expressed as mean ± SEM (n=3 biologically independent experiments, 3 donors). **P ≤ 0.01; ***P ≤ 0.001; ****P ≤ 0.0001 (one-way ANOVA with Dunnett correction for multiple comparisons. Statistical significance between mock and edited samples is depicted in the graph). **B**. Analysis of HbF and HbS by cation-exchange HPLC in RBCs derived from edited SCD HSPCs. We calculated the percentage of each Hb type over the total Hb tetramers. The EE ± SEM is indicated for each sample in the lower part of the panel. Data are expressed as mean ± SEM (n = 3 biologically independent experiments, 3 donors). *P ≤ 0.05; **P ≤ 0.01; ***P ≤ 0.001; ****P ≤ 0.0001 (two-way ANOVA with Dunnett correction for multiple comparisons. Statistical significance between mock and edited samples is depicted in the graph). **C-D**. C-G to T-A or/and A-T to G-C base-editing efficiency in the +58-kb (**C**) or +55-kb (**C**) regions calculated using EditR in samples subjected to Sanger sequencing in BFU-E colonies derived from SCD HSPCs edited at either the +58-kb or the +55-kb region, or simultaneously edited at both the +58-kb and the +55-kb regions. Data are expressed as mean (n = 2 biologically independent experiments, 2 donors). **E-F**. Frequency of InDels in the +58-kb (**E**) or +55-kb (**F**) regions measured by TIDE analysis, in samples subjected to Sanger sequencing in pools of BFU-E colonies differentiated from SCD HSPCs edited at either the +58-kb or the +55-kb regions, or simultaneously edited at both the +58-kb and the +55-kb regions. Data are expressed as mean ± SEM (n = 2 biologically independent experiments, 2 donors). **G**. Frequency of the 3.2-kb deletion/inversion, measured by ddPCR, in BFU-E colonies simultaneously edited at the +58-kb and +55-kb regions. Data are expressed as mean ± SEM (n = 2 biologically independent experiments, 2 donors). **H-I**. C-G to T-A or/and A-T to G-C base-editing efficiency in the +58-kb (**H**) or +55-kb (**I**) regions calculated using EditR in samples subjected to Sanger sequencing in CFU-GM colonies derived from SCD HSPCs edited at either the +58-kb or the +55-kb region, or simultaneously edited at both the +58-kb and the +55-kb regions. Data are expressed as mean (n = 2 biologically independent experiments, 2 donors). **J-K**. Frequency of InDels in the +58-kb (**J**) or +55-kb (**K**) regions measured by TIDE analysis, in samples subjected to Sanger sequencing in pools of CFU-GM colonies, differentiated from SCD HSPCs edited at either the +58-kb or the +55-kb regions, or simultaneously edited at both the +58-kb and the +55-kb regions. Data are expressed as mean ± SEM (n = 2 biologically independent experiments, 2 donors). **L.** Frequency of the 3.2-kb deletion/inversion, measured by ddPCR, in CFU-GM colonies simultaneously edited at the +58-kb and +55-kb regions. Data are expressed as mean ± SEM (n = 2 biologically independent experiments, 2 donors). *P ≤ 0.05; (two-way ANOVA with Sidak correction for multiple comparisons. Comparison of mock vs edited samples). **M.** Analysis of HbF and HbS by cation-exchange HPLC in pools of BFU-E colonies derived from edited SCD HSPCs. We calculated the percentage of each Hb type over the total Hb tetramers. Data are expressed as mean ± SEM (n = 2 biologically independent experiments, 2 donors).

**Supplementary Figure 4.**
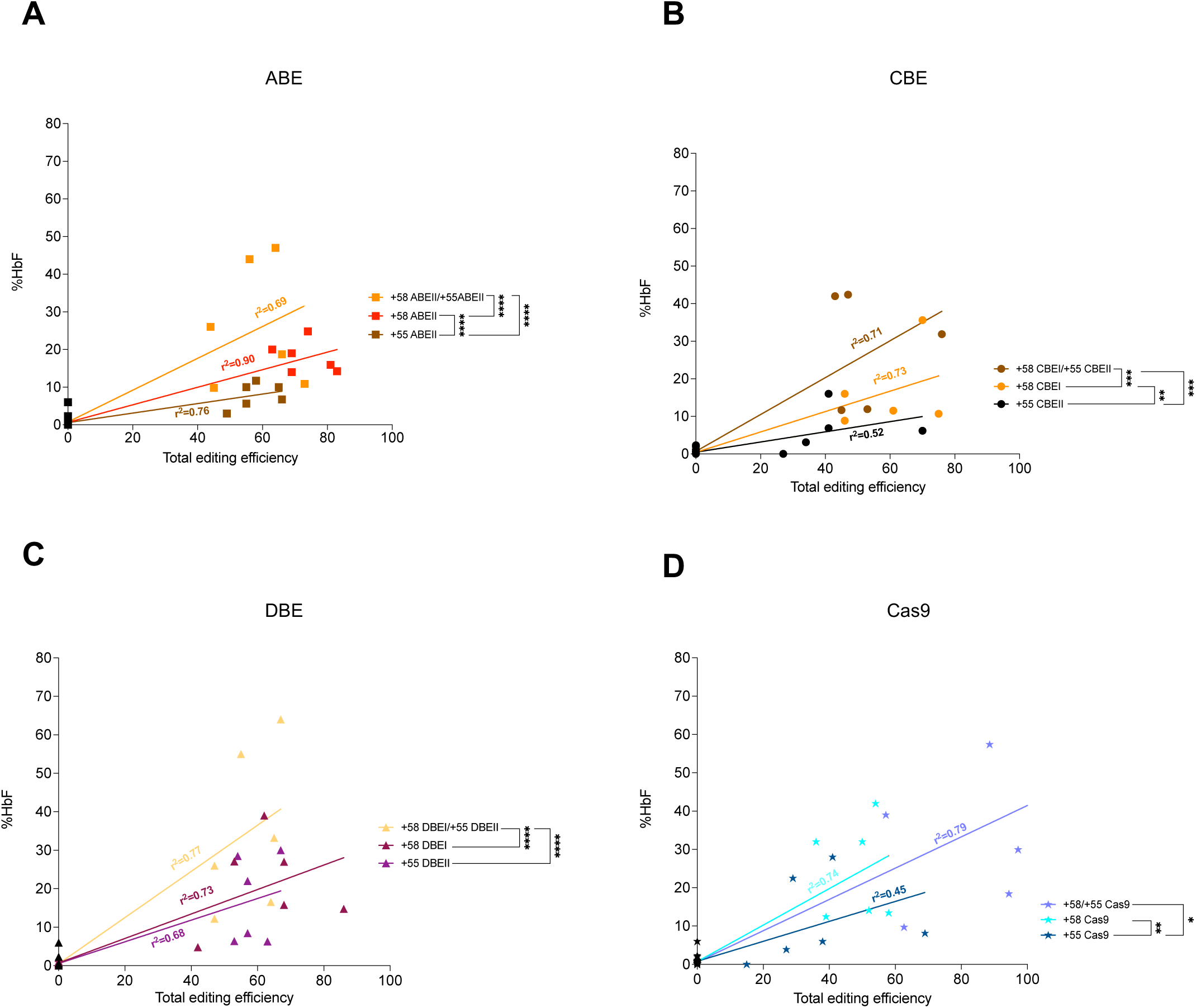
Correlation between HbF expression and editing efficiency obtained with ABE, CBE, DBE and Cas9 nuclease. Correlation between HbF expression and editing efficiency in erythroid cells derived from SCD HSPCs edited with either ABE (**A**), CBE (**B**), DBE (**C**) or Cas9-nuclease (**D**) (erythroblasts and pools of BFU-E; n = 3 biological independent experiments, 3 donors). HbF expression was measured by cation-exchange HPLC and calculated over the total Hb tetramers. Total editing efficiency was calculated by adding the base editing and Cas9 editing efficiency (determined by Sanger sequencing at the individual sites) to the frequency of the 3.2-kb deletion/inversion detected by ddPCR. Base-editing efficiency was calculated using EditR and Cas9-editing efficiency (InDels) was calculated using TIDE in samples subjected to Sanger sequencing. \**P* ≤ 0.05; \*\**P* ≤ 0.01; \*\*\**P* ≤ 0.001; \*\*\*\**P* ≤ 0.0001 (Multiple t test).

**Supplementary Figure 5.**
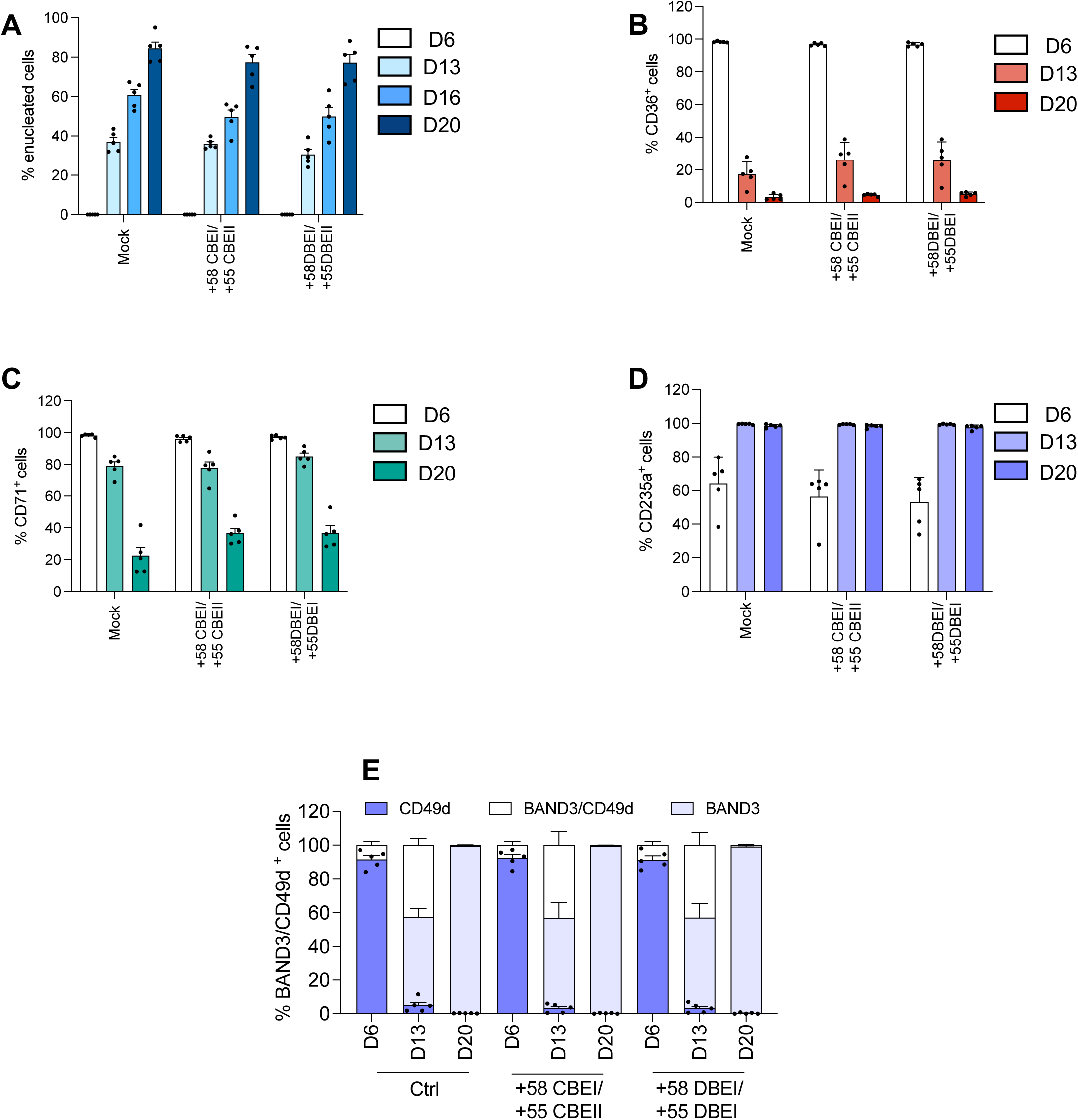
Erythroid differentiation of SCD HSPCs upon multiplex base editing of the +58-kb and +55-kb regions. **A.** Frequency of enucleated cells at day 6, 13, 16 and 20 of erythroid differentiation, as measured by flow cytometry analysis of DRAQ5 nuclear staining in control and edited samples. Data are expressed as mean ± SEM (n = 1 biologically independent experiment, 5 donors). **B-D**. Frequency of CD36^+^ (**B**), CD71^+^ (**C**) and CD235a^+^ (**D**) cells at day 6, 13 and 20 of erythroid differentiation, as measured by flow cytometry analysis of CD36, CD71 and CD235a erythroid markers. Data are expressed as mean ± SEM (n = 1 biologically independent experiment, 5 donors). **E**. Frequency of CD49d^+^, BAND3^+^ and CD49d^+^/BAND3^+^ in 7AAD^−^/CD235a^+^ cells at day 6, 13 and 20 of erythroid differentiation, as measured by flow cytometry analysis of CD49d and BAND3 erythroid markers. Data are expressed as mean ± SEM (n = 1 biologically independent experiment, 5 donors).

**Supplementary Figure 6.**
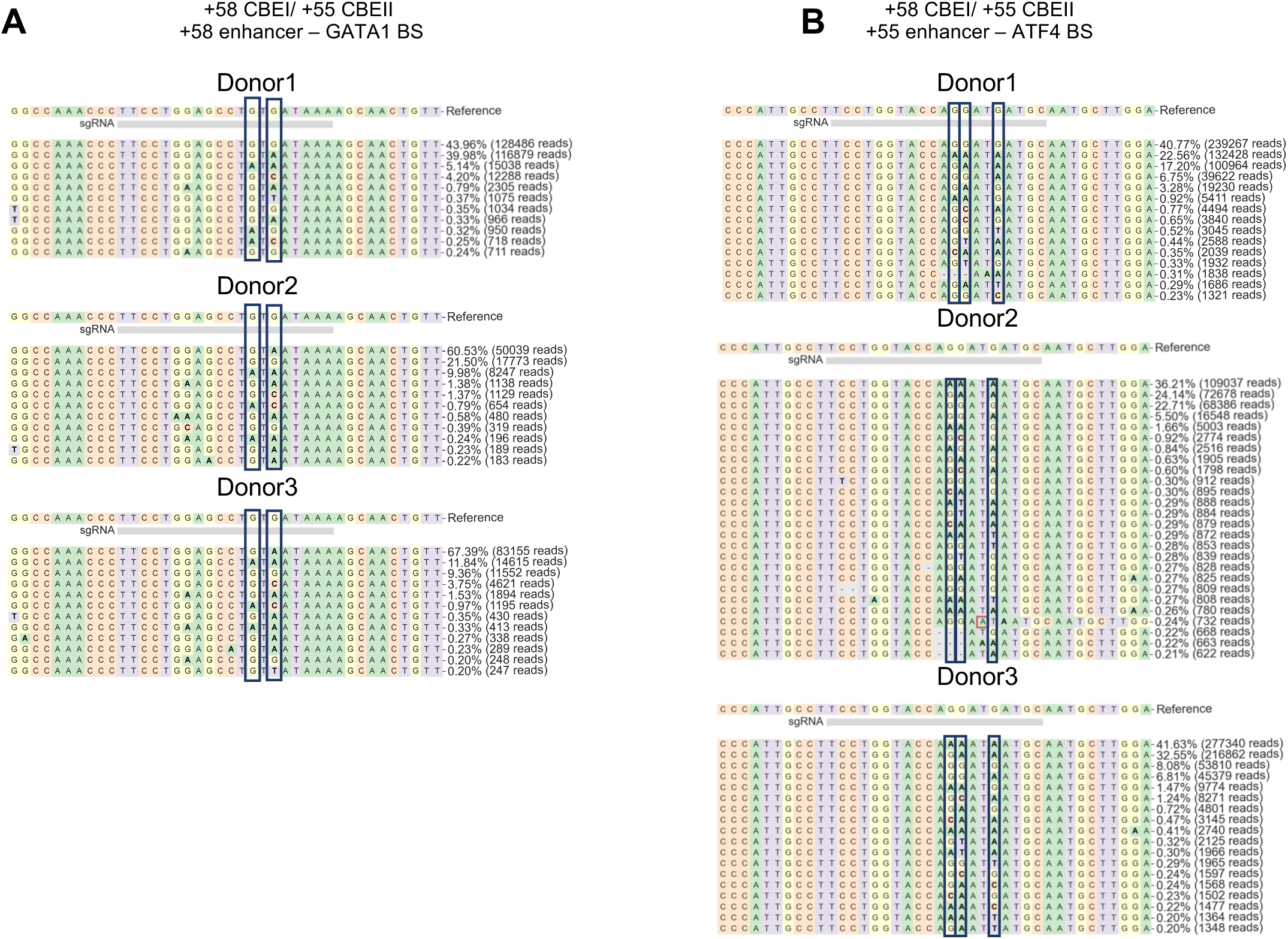
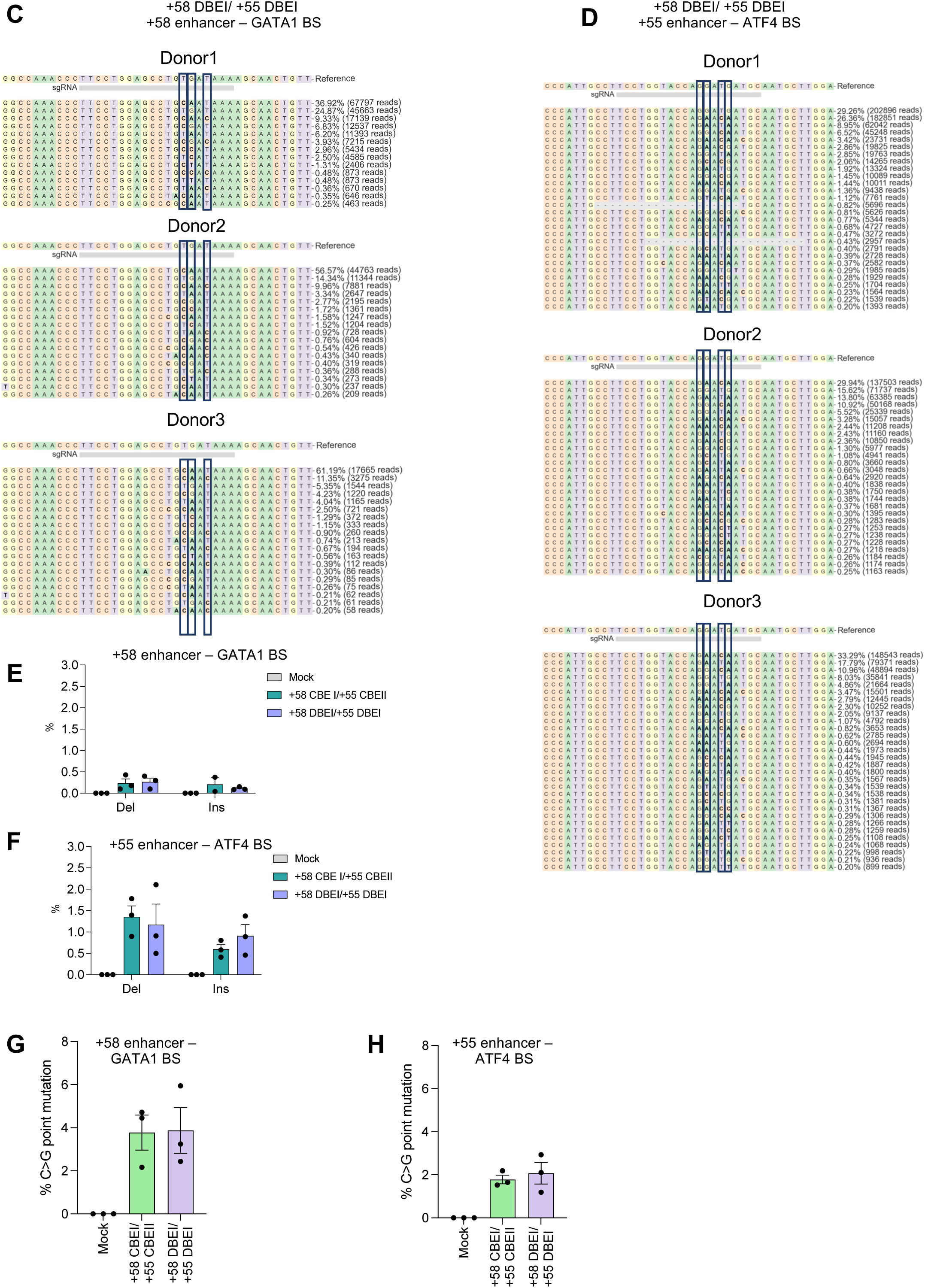
NGS analysis of on-target editing in erythroid cells derived from SCD HSPCs upon multiplex base editing of the +58-kb and +55-kb regions. **A-D.** Frequency and sequence of modified and unmodified alleles in edited SCD samples, for +58 CBEI/ +55 CBEII (**A** and **B**) and +58 DBEI/ +55 DBEI (**C** and **D**) profiles at the +58-kb (**A** and **C**) and +55-kb (**B** and **D**) regions, as measured by targeted NGS. Target base positions are highlighted with a blue box. Red box indicates inserted bases. Grey squares indicate deletions. (n = 3 biologically independent experiments, 3 donors). **E-F.** Deletions and Insertions frequency in edited SCD samples for +58 CBEI/ +55 CBEII and +58 DBEI/ +55 DBEI profiles at the +58-kb (**E**) and +55-kb (**F**) regions, as measured by targeted NGS. Data are expressed as mean ± SEM (n = 3 biologically independent experiments, 3 donors). **G-H.** C-G to G-C base-editing frequency in the +58-kb (**G**) or +55-kb (**H**) regions as measured by targeted NGS in SCD HSPCs simultaneously edited at the +58-kb and the +55-kb region. Data are expressed as mean ± SEM (n = 3 biologically independent experiments, 3 donors).

**Supplementary Figure 7.**
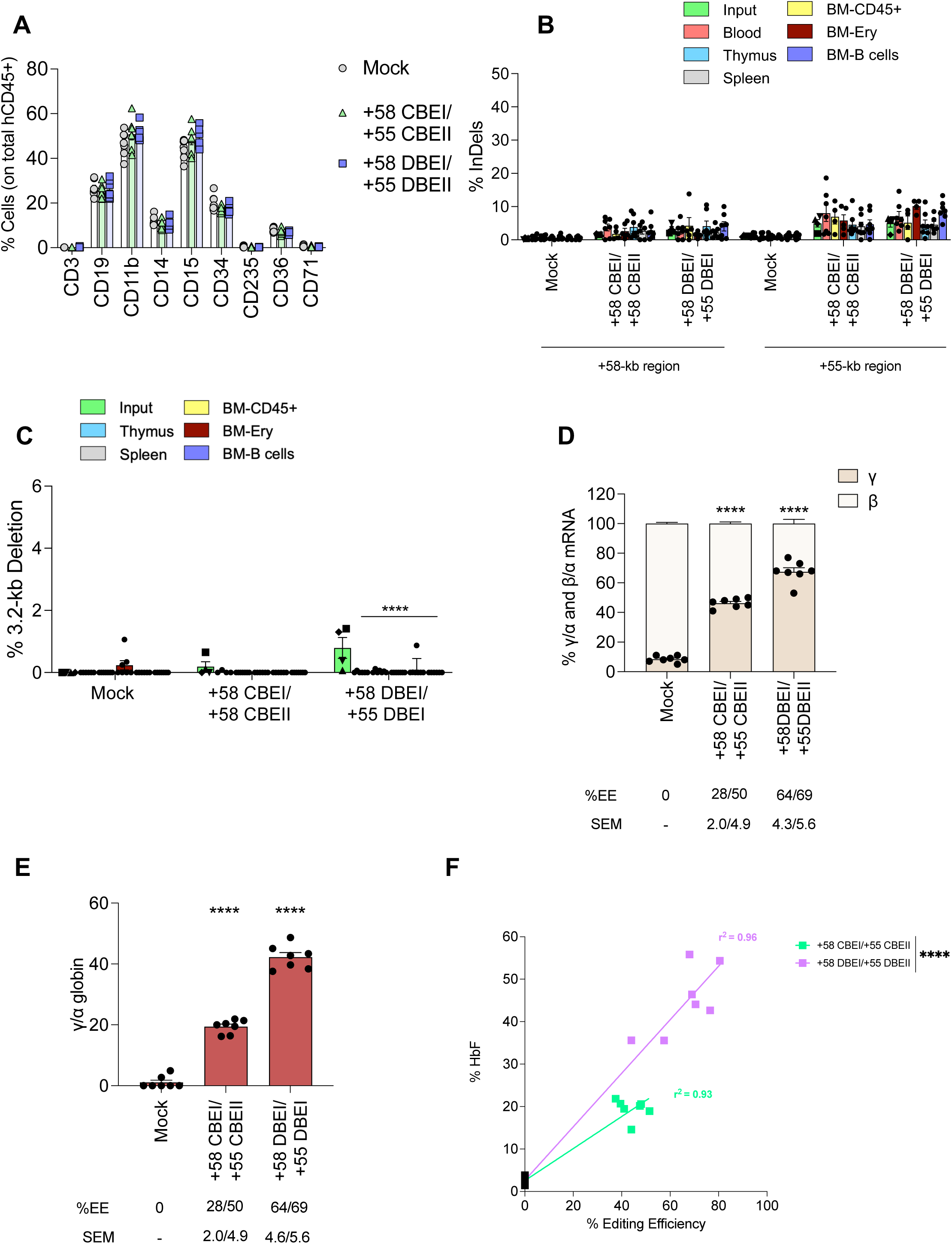
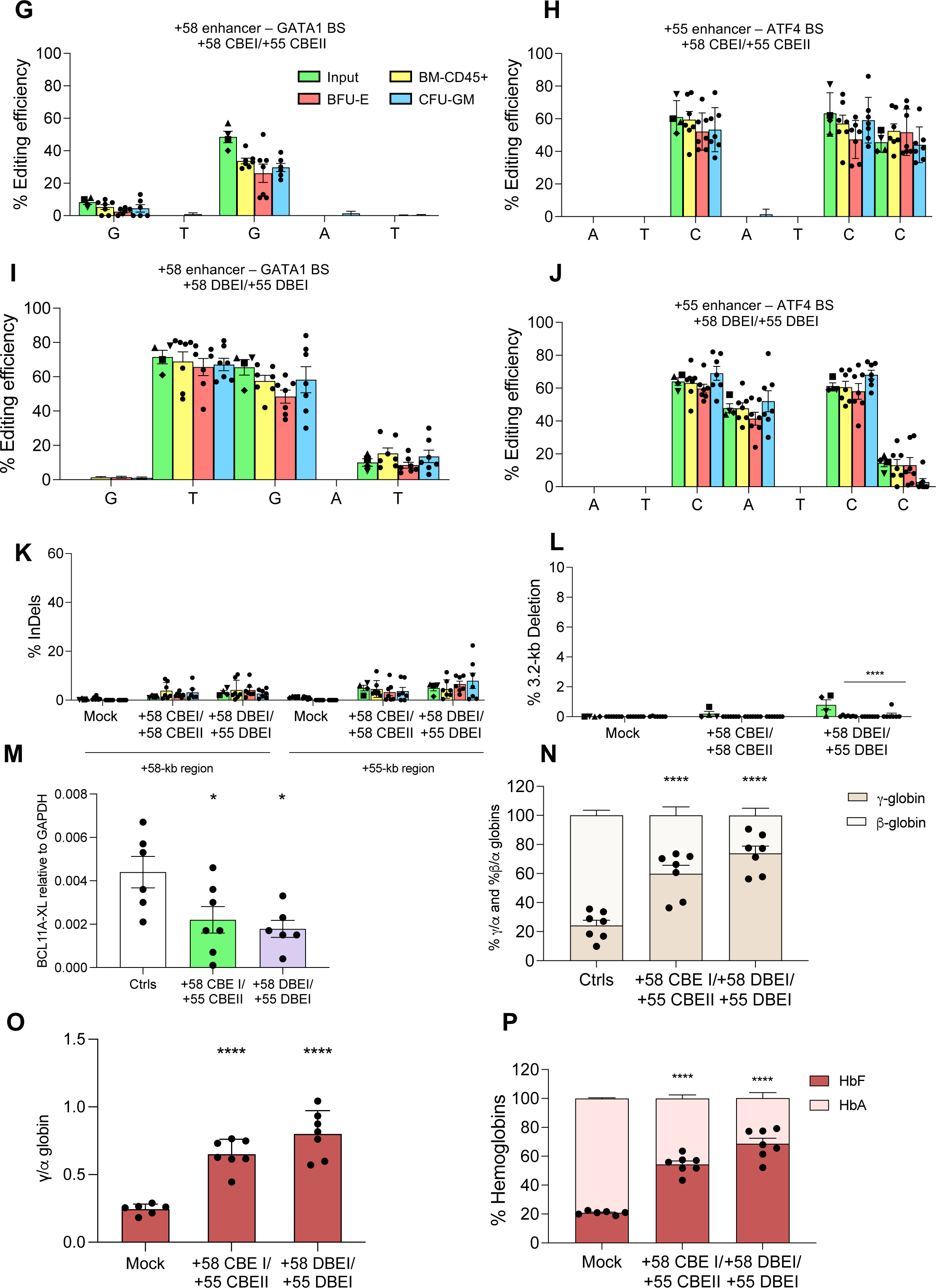
Simultaneous base editing of *BCL11A* enhancers in repopulating HSCs and erythroid and granulo-monocytic progenitors derived from engrafting HSPCs. **A.** Frequency of human T (CD3) and B (CD19) lymphoid, myeloid (CD11b, CD14, and CD15), HSPC (CD34), and erythroid (CD235a, CD36 and CD71) cells in BM in mice transplanted with control and edited HSPCs, 16/17 weeks after the transplantation (n = 7 mice per condition). Each data point represents an individual mouse. Data are expressed as mean ± SEM. (two-way ANOVA; not significant). **B.** Frequency of InDels calculated by TIDE for the +58 CBEI/+55 CBEII and +58 DBEI/+55 DBEI profiles at the +58-kb and +55-kb regions in the input, blood-, BM-, thymus-, and spleen-derived human samples subjected to Sanger sequencing. BM samples include CD45^+^ cells, B cells and erythroid cells (Ery). Data are expressed as mean ± SEM [n = 1 biologically independent experiment (Input), n = 4 to 7 mice per group]. Each data point represents an individual mouse (two-way ANOVA; not significant). The frequency of Indels in the input was calculated in cells cultured in the HSPC medium (▪), in liquid erythroid cultures (▲), and in pools of BFU-E (◆) and CFU-GM (▾). **C.** Frequency of the 3.2-kb deletion, measured by ddPCR, for samples simultaneously edited at the +58-kb and +55-kb regions for the input, BM-, thymus-, and spleen-derived human samples. BM samples include CD45^+^ cells, B cells and erythroid cells (Ery). Data are expressed as mean ± SEM [n = 1 biologically independent experiment (Input), n = 7 mice per group]. Each data point represents an individual mouse. \*\*\*\**P* ≤ 0.0001 between the input and the different human hematopoietic populations derived from engrafted HSCs (two-way ANOVA with Tukey’s correction for multiple comparisons). The frequency of 3.2-kb deletion in the input was calculated in cells cultured in the HSPC medium (▪), in liquid erythroid cultures (▲), and in pools of BFU-E (◆) and CFU-GM (▼). **D.** RT-qPCR analysis of γ (^G^γ + ^A^γ)- and β-globin mRNA in BM sorted human CD235a^+^ erythroid cells in mice transplanted with control and edited HSPCs, 16/17 weeks post-transplantation. γ- and β^S^-globin mRNA expression was normalized to α-globin mRNA and expressed as a percentage of the γ-+ β^S^-globin mRNA. The EE ± SEM is indicated for each sample in the lower part of the panel. Each data point represents an individual mouse. Data are expressed as mean ± SEM (n = 7 mice per group). \*\*\*\**P* ≤ 0.0001 (two-way ANOVA with Dunnett correction for multiple comparisons. Comparison mock vs edited samples). **E.** Expression of γ (^G^γ + ^A^γ)-globin chains measured by RP-HPLC in BM sorted human CD235a^+^ erythroid cells in mice transplanted with control and edited HSPCs, 16/17 weeks post-transplantation. γ-globin expression was normalized to α-globin. The EE ± SEM is indicated for each sample in the lower part of the panel. Each data point represents an individual mouse. Data are expressed as mean ± SEM (n = 7 mice per group). \*\*\*\**P* ≤ 0.0001 (one-way ANOVA with Dunnett correction for multiple comparisons. Comparison mock vs edited samples). **F.** Correlation between HbF expression and editing efficiency in human CD235a^+^ BM-sorted erythroid cells obtained from mice transplanted with control and edited HSPCs. HbF was measured by cation-exchange HPLC and calculated over the total Hb tetramers. Base-editing efficiency was calculated using EditR software. ****P ≤ 0.0001 (Multiple t test). **G-J**. C-G to T-A or/and A-T to G-C base-editing efficiency, calculated using EditR for the +58 CBEI/ +55 CBEII (**G** and **H**) and +58 DBEI/ +55 DBEI (**I** and **J**) profiles at the +58-kb (**G** and **I**) and +55-kb (**H** and **J**) regions in the input, BM CD45^+^ cells and pooled BFU-E and CFU-GM derived from BM CD45^+^ cells and subjected to Sanger sequencing. Data are expressed as mean ± SEM [n = 1 biologically independent experiment (Input), n = 7 mice per group]. Each data point represents an individual mouse. The frequency of base editing in the input was calculated in cells cultured in the HSPC medium (▪), in liquid erythroid cultures (▲) and in pools of BFU-E (◆) and CFU-GM (▾). **L.** Frequency of InDels calculated by TIDE for the +58 CBEI/+55 CBEII and +58 DBEI/+55 DBEI profiles at the +58-kb and +55-kb regions in the input, BM CD45^+^ cells, and pooled BFU-E and CFU-GM derived from BM CD45^+^ cells and subjected to Sanger sequencing. Data are expressed as mean ± SEM [n = 1 biologically independent experiment (Input), n = 7 mice per group]. Each data point represents an individual mouse (two-way ANOVA; not significant). The frequency of Indels in the input was calculated in cells cultured in the HSPC medium (▪), in liquid erythroid cultures (▴), and in pools of BFU-E (◆) and CFU-GM (▾). **M.** Frequency of the 3.2-kb deletion, measured by ddPCR, for samples simultaneously edited at the +58-kb and +55-kb regions for the input, BM CD45^+^ cells, and pooled BFU-E and CFU-GM derived from BM CD45^+^ cells. Data are expressed as mean ± SEM [n = 1 biologically independent experiment (Input), n = 7 mice per group]. Each data point represents an individual mouse. \*\*\*\**P* ≤ 0.0001 between the input and the different human hematopoietic populations derived from engrafted HSCs (two-way ANOVA with Tukey’s correction for multiple comparisons). The frequency of 3.2-kb deletion in the input was calculated in cells cultured in the HSPC medium (▪), in liquid erythroid cultures (▲), and in pools of BFU-E (◆) and CFU-GM (▾). **N.** RT-qPCR analysis of *BCL11A-XL* expression in pooled BFU-E derived from BM CD45^+^ cells. *BCL11A-XL* mRNA expression was normalized to *GAPDH*. Data are expressed as mean ± SEM (n = 7 mice per group). Each data point represents an individual mouse. *P ≤ 0.05 (One-way ANOVA with Dunnett’s correction for multiple comparison. Comparison of mock vs edited samples). **O.** RT-qPCR analysis of γ (^G^γ + ^A^γ)- and β-globin mRNA in pooled BFU-E derived from BM CD45^+^ cells. γ- and β^−^globin mRNA expression was normalized to α-globin mRNA and expressed as a percentage of the γ- + β- globin mRNA. Each data point represents an individual mouse. Data are expressed as mean ± SEM (n = 7 mice per group). \*\*\*\**P* ≤ 0.0001 (two-way ANOVA with Dunnett) **P.** Expression of γ (^G^γ + ^A^γ)-globin chains measured by RP-HPLC in pooled BFU-E derived from BM CD45^+^ cells. γ-globin expression was normalized to α-globin. Each data point represents an individual mouse. Data are expressed as mean ± SEM (n = 7 mice per group). \*\*\*\**P* ≤ 0.0001 (one-way ANOVA with Dunnett correction for multiple comparisons. Comparison mock vs edited samples). **Q.** Analysis of HbF and HbS by CE-HPLC in pooled BFU-E derived from BM CD45^+^ cells. We calculated the percentage of each Hb type over the total Hb tetramers. Data are expressed as mean ± SEM (n = 7 mice per group). Each data point represents an individual mouse. ****P ≤ 0.0001 (two-way ANOVA with Dunnett correction for multiple comparisons. Comparison of mock vs edited samples).

## Acknowledgements

This work was supported by state funding from the French National Research Agency (Agence Nationale de la Recherche) as part of the Investissements d’Avenir program (ANR-10-IAHU-01) and by the European Research Council (865797 DITSB), the European Commission (HORIZON-RIA EDITSCD grant no. 101057659), the Fondation pour la Recherche Médicale [FRM PLP202110014595] and the COST (European Cooperation in Science and Technology (the COST Action Gene Editing for the treatment of Human Diseases, CA21113).

We thank Carine Giovannangeli for the production of Cas9 nuclease RNP and all the patients for their contribution to this work.

## Author contributions

L.F. designed, conducted experiments, analyzed the data, and wrote the paper. P.M., S.A., T.F., M.M., G.C., A.T., A.C., G.H., M.J. conducted experiments and analyzed data. O.R. analyzed NGS data. M.A. contributed to the design of the experimental strategy. P.A. conceived the study, designed, conducted experiments, analyzed the data, and wrote the paper. A.M. conceived the study, designed experiments, and wrote the paper.

